# Understanding the state-dependent impact of species correlated responses on community sensitivity to perturbations

**DOI:** 10.1101/2022.07.24.501326

**Authors:** Lucas P. Medeiros, Serguei Saavedra

## Abstract

Understanding how communities respond to perturbations requires us to consider not only changes in the abundance of individual species, but also correlated changes that can emerge through interspecific effects. However, our knowledge of this phenomenon is mostly constrained to populations at equilibrium, where interspecific effects are fixed. Here, we introduce a framework to disentangle the impact of species correlated responses on community sensitivity to perturbations when interspecific effects change over time due to non-equilibrium dynamics. We partition the volume expansion rate of perturbed abundances (community sensitivity) into contributions of individual species and of species correlations by converting the time-varying Jacobian matrix containing interspecific effects into a time-varying covariance matrix. Using population dynamics models, we demonstrate that species correlations change considerably across time and continuously alternate between reducing and having no impact on community sensitivity. Importantly, these alternating impacts depend on the abundance of particular species and can be detected even from noisy time series. We showcase our framework using two experimental predator-prey time series and find that the impact of species correlations is modulated by prey abundance—as theoretically expected. Our results provide new insights into how and when species interactions can dampen community sensitivity when abundances fluctuate over time.

## Introduction

Natural and human-driven perturbations such as fires (1), storms (2), pollution (3), and over-fishing (4) can alter the composition of ecological communities as well as the abundance of their constituent species. Thus, there is an urgent need to understand the capacity of communities to retain their biodiversity and functioning in the face of rapidly increasing environmental perturbations (5, 6). It is well-known that the way communities respond to perturbations (e.g., recovery, constancy, sensitivity) cannot be explained solely by the sum of responses of isolated species, but depends crucially on the interactions among these species (7–10). Indeed, because all species are not equal and species may affect each other’s response to perturbations, scaling up from individual species through their interactions to understand the response of an entire community to perturbations remains an important challenge.

A fundamental consequence of species interactions is that changes in the abundance of a given species following a perturbation may cascade and shift the abundance of other species, creating correlations in how species respond to perturbations. Such correlated species responses include, for example, shifts in preys (or resources) immediately following changes in the abundance of predators (or consumers) (11–13); and shifts in competitors immediately following changes in environmental conditions (9, 14, 15). Importantly, such correlations can impact the response of aggregate community properties following perturbations. For instance, when measuring community response as constancy of total abundance over time, opposite changes in the abundance of different species in the face of perturbations (i.e., negative correlations) will increase community constancy—a phenomenon known as compensatory dynamics (9, 14–16). Given that many ecosystem services (e.g., carbon sequestration, food provisioning, pollination) depend on total abundance, compensatory dynamics has been invoked as a central mechanism stabilizing the provision of these services (5, 16).

Despite important progress in assessing how species correlated responses affect whole-community response to perturbations, the fact that this effect appears to be context-dependent has posed big challenges to its understanding. Key examples come from food web studies, where the effect of removing or adding a predator on other species in the community critically depends on the sign and strength of species interactions (17–19). Thus, depending on species initial abundances and their interactions, two species may show a positive, negative, or null correlation in their response to a perturbation. Such context-dependency becomes clearer when we consider that, in the majority of natural communities, the instantaneous effect of each species on the growth rate of other species in the community (hereafter interspecific effects) shifts over time due to changes in community state (i.e., distribution of species abundances) (20–22). That is, even when the sign and strength of the per-capita effect of a species on the per-capita growth rate of another species is fixed (e.g., fixed pairwise interaction parameters in a population dynamics model), interspecific effects (i.e., elements of the Jacobian matrix of a model) will change when species abundances change (23, 24). The fact that interspecific effects are state-dependent is particularly important when populations do not settle down to an equilibrium (e.g., cyclic or chaotic dynamics), which is the case for many communities (20, 21, 25–28). What is currently unknown is how such interspecific effects affect species correlated responses to perturbations and, in turn, how such correlations impact whole-community response under non-equilibrium dynamics.

Moving beyond the typical assumption of populations at equilibrium, recent work has begun to elucidate how communities respond to perturbations under non-equilibrium dynamics (20, 29). For example, it has been shown that, not only interspecific effects, but also the community’s response to perturbations changes over time when dynamics are out of equilibrium (20, 29). In particular, a community under non-equilibrium dynamics may have a time-varying sensitivity to changes in environmental conditions (29, 30). In addition to whole-community response, the sensitivity of individual species to perturbations has also been shown to change over time, implying that the identity of the most sensitive species also depends on community state (31). In spite of this recent progress, the current frameworks to quantify response to perturbations at the community level (20, 29) and at the species level (31) under non-equilibrium dynamics have remained disconnected. Yet, investigating potential time-varying correlations in how species respond to perturbations may provide the key to understand the links between responses to perturbations at these two levels of biological organization.

Here, we introduce a theoretical framework to assess the time-varying impact of species correlated responses (hereafter species correlations) on the sensitivity of a community to perturbations under non-equilibrium dynamics. In what follows, we first present our framework that consists of partitioning community sensitivity—measured as the volume expansion rate of perturbed abundances—into contributions of individual species and of species correlations by converting the time-varying Jacobian matrix into a time-varying covariance matrix. Then, we illustrate our framework using synthetic time series generated from population dynamics models and show how we can identify community states where species correlations have either a weak or strong impact on community sensitivity. Lastly, we apply our framework to two experimental predator-prey communities and confirm that prey abundance is an important factor determining whether or not species correlations can dampen community sensitivity to perturbations across time.

## Results

### Species correlated responses and their impact on community sensitivity

In general, the population dynamics of a community with *S* species can be written as: 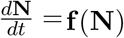, where **N** = [*N*_1_, …, *N*_*S*_]^┬^ is the vector of species abundances and **f** = (*f*_1_, …, *f*_*S*_) (*f*_*i*_: ℝ^*S*^ →ℝ) is a set of generic functions describing the growth rates of abundances (32). As an example, consider the population dynamics model of a community with one predator and one prey (*Materials and Methods*). Under certain parameter values, this model generates a limit cycle (33), which we use to illustrate the state-dependent impact of species correlations on whole-community sensitivity to perturbations (Fig. 1). At any given state (i.e., any **N** along the cycle), this community may be affected by a pulse perturbation **p** = [*p*_1_, …, *p*_*S*_]^┬^ that changes **N** into Ñ (i.e., Ñ = **N** + **p**) (34). The vector Ñ would then change in time according to **f**. For instance, consider a perturbation that decreases the abundance of the predator (**p** = [−7, 0]^┬^, red arrow in Figs. 1a, b) and an opposite perturbation with the same magnitude that increases it (**p** = [7, 0]^┬^, blue arrow in Figs. 1a, b). Figure 1a shows the impact of these two perturbations on the prey abundance (Δ*N*_2_) at time *t* = *t*_1_ and after *k* time steps. The figure shows that although one perturbation decreased the predator abundance and the other increased it, the impacts on the prey abundance (red Δ*N*_2_ vs blue Δ*N*_2_ in Fig. 1a) are similar to each other. Thus, knowing whether the perturbation decreases or increases the predator abundance gives little information about the amount of change in the prey abundance (i.e., species responses to perturbations at time *t*_1_ are uncorrelated). Interestingly, however, the same perturbations applied at time *t* = *t*_2_ can trigger a completely different outcome (Fig. 1b). As can be seen in Figure 1b, the impact on the prey abundance is much larger when the predator abundance is decreased (red Δ*N*_2_ in Fig. 1b) than when it is increased (blue Δ*N*_2_ in Fig. 1b). In this case, knowing how the perturbation impacts the predator abundance gives information about how much change we are likely to see in the prey abundance (i.e., species responses to perturbations at time *t*_2_ are correlated).

**Figure 1.**
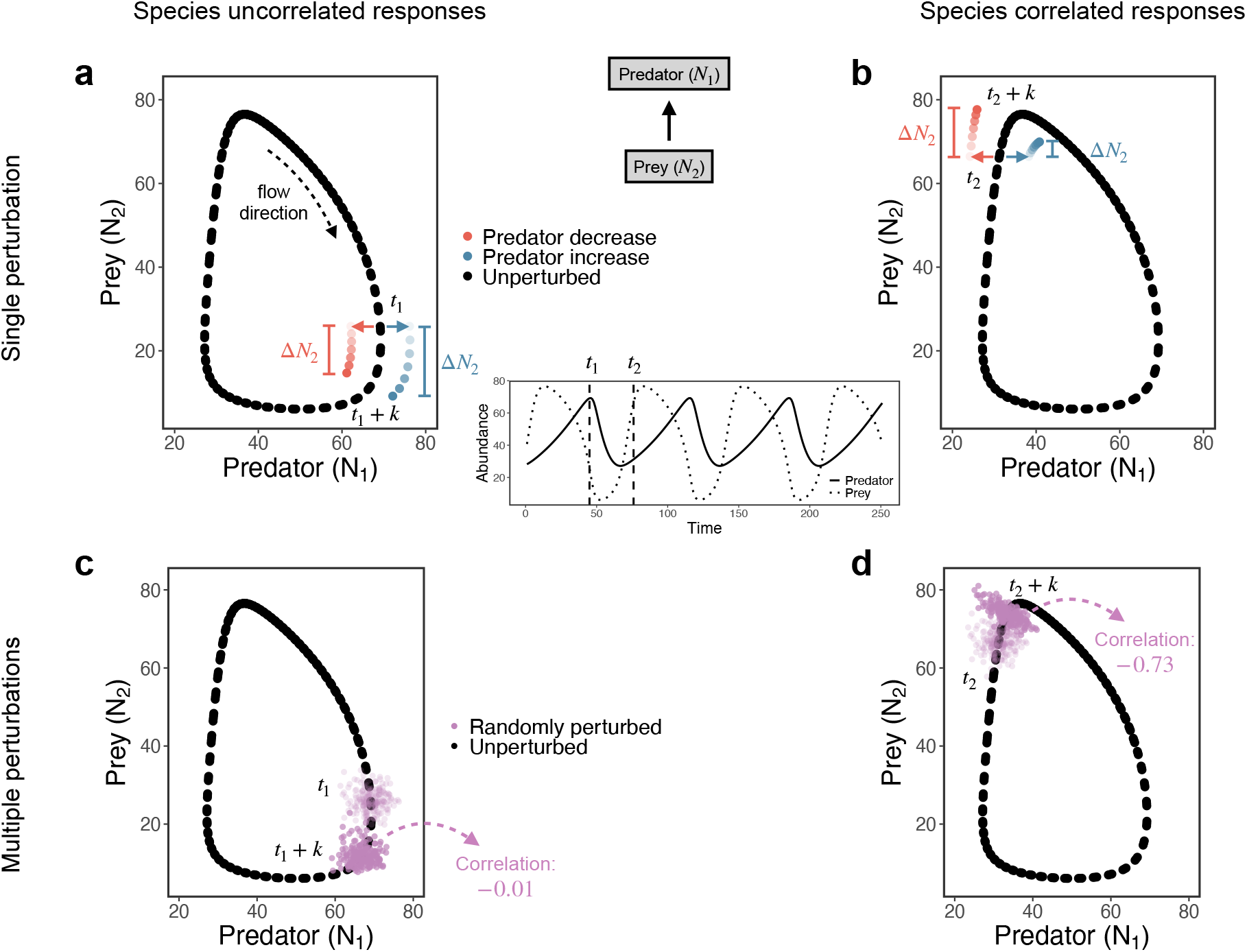
Correlations in how species respond to perturbations depend on community state under non-equilibrium dynamics. **a**-**b**, Limit cycle of a 2-species predator-prey model (black points; equation [2]) with two pulse perturbations (**p** = [−7, 0]^⊤^, red arrow and **p** = [7, 0]^⊤^, blue arrow) affecting the abundance of the predator (*N*_1_) at two different points in time (*t* = *t*_1_ and *t* = *t*_2_). The central panel shows the abundance time series (solid line: predator; dotted line: prey) and the two time points as vertical dashed lines. **a**, At time *t*_1_, the effect of both perturbations on prey abundance after *k* = 3 time steps (Δ*N*_2_) is similar (Δ*N*_2_ = −11.1 when predator decreases, in red; Δ*N*_2_ = −16.7 when predator increases, in blue). **b**, In contrast, at time *t*_2_, one perturbation has a much larger effect on prey abundance than the other (Δ*N*_2_ = 11.2 when predator decreases, in red; Δ*N*_2_ = 3.5 when predator increases, in blue). **c**-**d**, Same limit cycle as in **a**-**b**, but showing the outcome of multiple random perturbations (*p*_*i*_ ∼𝒩 (*µ* = 0, *σ*^2^ = 9)) that change **N** (black point) into **Ñ** (light purple points) at times *t*_1_ (**c**) and *t*_2_ (**d**). **c**, At time *t*_1_ + *k*, there is no correlation between the perturbed abundances of the predator (*Ñ*_1_) and of the prey (*Ñ*_2_) (dark purple points; correlation: -0.01). **d**, In contrast, at time *t*_2_ + *k*, there is a strong correlation between the perturbed abundances of the predator and of the prey (dark purple points; correlation: -0.73).

Although the previous example is based on only two perturbations (decreasing or increasing the predator abundance), we can observe the same outcomes when considering several random perturbations **p** around **N** (Figs. 1c, d). It is worth noting that here we focus on random perturbations as we typically have no a priori information about how external perturbations will change natural communities. Each light purple point in Figures 1c, d represents a vector of perturbed abundances (**Ñ**; total of 200 vectors), where *p*_*i*_ ∼ 𝒩 (*µ* = 0, *σ*^2^ = 9). Figure 1c shows the distribution of perturbed abundances at *t* = *t*_1_ (light purple points) and after *k* time steps (dark purple points). As expected from Figure 1a, the correlation between *Ñ*_1_ and *Ñ*_2_ (computed from all 200 perturbed abundances) at *t*_1_ + *k* is almost zero (*ρ* = 0.01; Fig. 1c). Similarly, Figure 1d shows the distribution of perturbed abundances at *t* = *t*_2_ (light purple points) and after *k* time steps (dark purple points). Again, as expected from Figure 1b, the correlation between *Ñ*_1_ and *Ñ*_2_ at *t*_2_ + *k* is strong and negative (*ρ* = −0.76; Fig. 1d). This numerical exercise illustrates the main question we address in this study: how the absence (Fig. 1c) or presence (Fig. 1d) of species correlations at different community states impact whole-community sensitivity to perturbations?

### Decomposing community sensitivity to perturbations

Under equilibrium dynamics (i.e., **N** = **N**^∗^ with **f** (**N**^∗^) = **0**), the response of a community to pulse perturbations has been studied using several different indicators such as recovery rate (35) and reactivity (36). Nevertheless, under non-equilibrium dynamics it is necessary to establish a measure of community response that does not depend on a return to an equilibrium point. Therefore, here we focus on a measure known as the volume expansion rate, which is related to how much change we expect to see in **N** under perturbations—that is, the community sensitivity to perturbations (Fig. 2a) (29, 30). Note that the inverse of this measure, which has been termed volume contraction rate, can be interpreted as the community resistance to perturbations (30). Although this measure was originally used to measure sensitivity to perturbations on the governing dynamical laws of a community (29), we prove that this measure is also informative of perturbations on species abundances (*Materials and Methods*). Below, we derive our framework of partitioning community sensitivity into contributions of individual species and of species correlations.

**Figure 2.**
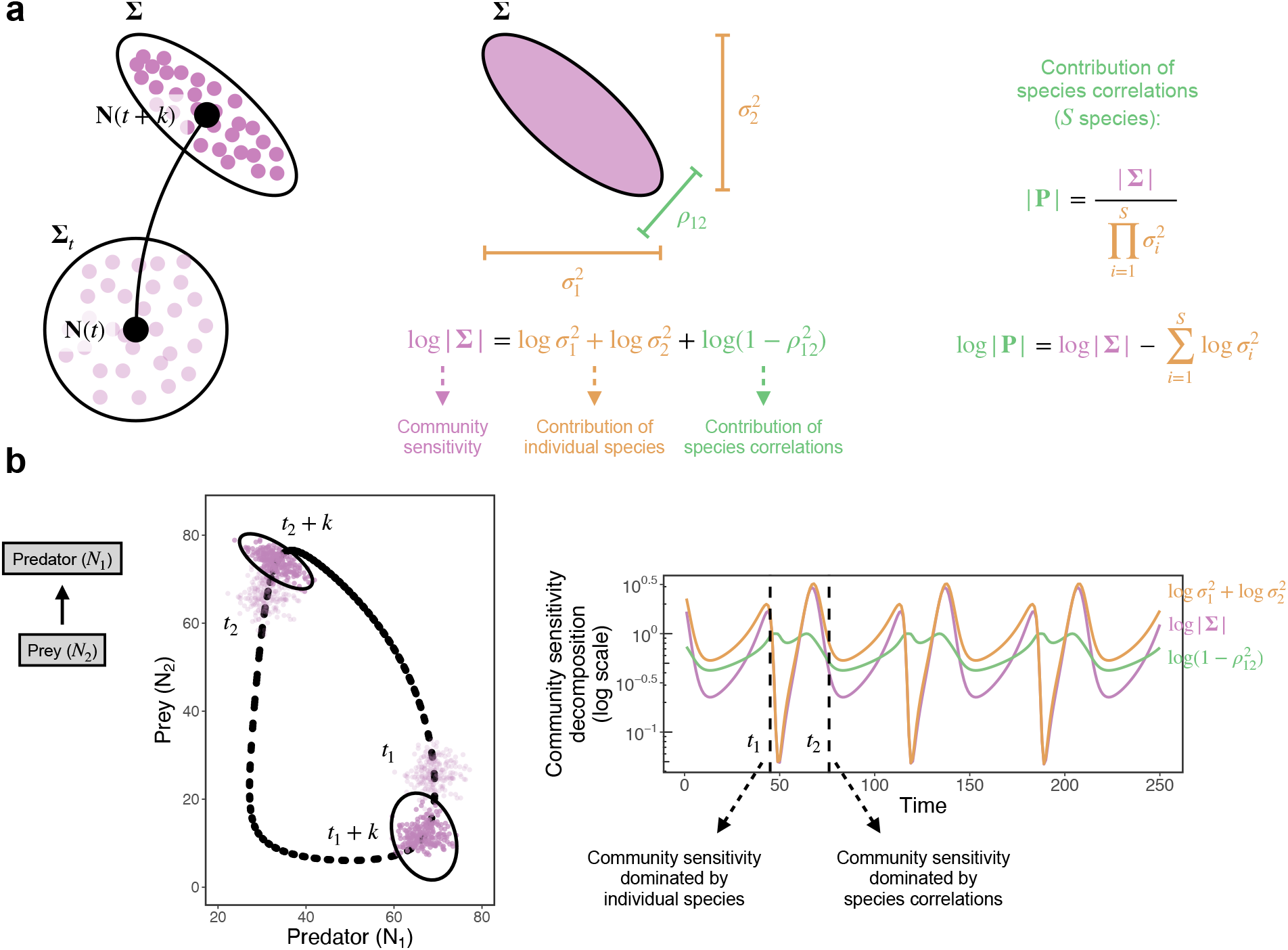
Decomposing community sensitivity into contributions of individual species and of species correlated responses. **a** (left), Diagram of a non-equilibrium trajectory showing how perturbed abundances at time *t* (light purple points) generated using covariance matrix **Σ**_*t*_ (black circle) can be described at time *t* + *k* (dark purple points) by covariance matrix **Σ** (black ellipse). **a** (center), Decomposition of the community sensitivity to perturbations (volume expansion rate, log |**Σ**|) into contributions of individual species (variances in 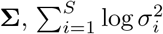) and of species correlations (determinant of correlation matrix, log |**P**|), which is shown for 2 species. **a** (right), With *S* species, we can compute the relative importance of species correlations as 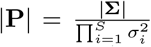, where |**P**| → 0 indicates that community sensitivity is minimized due to species correlations. **b** (left), Same predator-prey limit cycle (black points; equation [2]) and time points (*t* = *t*_1_ and *t* = *t*_2_) as in Fig. 1 showing that **Σ** (black ellipses) indeed captures the lack of correlation of perturbed abundances at time *t*_1_ + *k* and the strong correlation at time *t*_2_ + *k* (*k* = 3). **b** (right), Community sensitivity decomposition at each point in time for the 2-species model. Whereas community sensitivity (purple line) is dominated by the contribution of individual species (orange line) at time *t*_1_ (|**P**| ≈ 1), it is dominated by the contribution of species correlations (green line) at time *t*_2_ (|**P**| ≪ 1).

As mentioned before, because we typically have no information about the effect of external perturbations, here we assume that pulse perturbations at an arbitrary time *t* (**p**(*t*)) follow a distribution with mean vector ***µ***_*t*_ and covariance matrix **Σ**_*t*_ (Fig. 2a; *Materials and Methods*). Using the linearized dynamics of **p**, we can then obtain the covariance matrix of **p**(*t* + *k*) as: 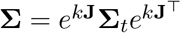 (*Materials and Methods, Supporting Information Section 1*). Thus, **Σ** is obtained via a transformation of the initial covariance matrix (**Σ**_*t*_) due to interspecific effects present in the Jacobian matrix **J** (Fig. 2a). We assume that **J** is almost constant from time *t* to *t* + *k*, which is a reasonable assumption whenever *k* or 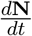 are small (31). Note, however, that **J** can change across a non-equilibrium attractor because it depends on the community state **N** (i.e., it is state-dependent). Also note that, in addition to knowing **J**, knowledge of **Σ**_*t*_ and *k* is required to compute **Σ**. The accuracy of **Σ** in describing the distribution of perturbed abundances has been verified under equilibrium dynamics (35), whereas here we perform simulations to confirm this accuracy under non-equilibrium dynamics (*Supporting Information Section 2* ; Fig. S1).

Without loss of generality, we can assume that **p**(*t*) and, therefore, **p**(*t*+*k*) follows a multivariate normal distribution. In this case, the covariance matrices **Σ**_*t*_ and **Σ** can be visualized as 95% confidence *S*-dimensional ellipses surrounding perturbed abundances (Fig. 2a). Importantly, the determinants |**Σ**_*t*_| and |**Σ**| are proportional to the volume of such ellipses (37–39), and represent the overall change in species abundances following random pulse perturbations (i.e., community sensitivity). Alternatively, the change of an infinitesimal volume around **N** (i.e., volume expansion rate) is given by the divergence of the vector field **f** : 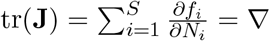. **f** (**N**) (29, 40). We show that |**Σ**| is equivalent to tr(**J**) via the following expression: log |**Σ**| = log (*v*) + 2*k*tr(**J**), where |**Σ**_*t*_| = *v* is the arbitrary volume of initial perturbations (*Materials and Methods*). Thus, we define log |**Σ**| as our measure of state-dependent community sensitivity to perturbations and conclude that, for a given value of *v* and *k*, a larger log |**Σ**| implies a larger tr(**J**). Note that abundance changes have to be interpreted in terms of their units and abundances are typically normalized when working with empirical data (*Materials and Methods*).

Because the determinant of any covariance matrix can be written as the product of variances 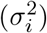 times the determinant of the correlation matrix **P** (*Supporting Information Section 3*), we can decompose the community sensitivity (log |**Σ**|) into contributions of individual species and of species correlations:

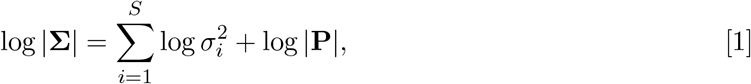

where the determinant of **P** (|**P**|) depends on all correlations *ρ*_*ij*_ between species *i* and *j* (*i, j* = 1, …, *S*). Note that *ρ*_*ij*_ is calculated by dividing the covariance between species *i* and *j* (i.e., *ij*th element of **Σ**) by *σ*_*i*_*σ*_*j*_. Therefore, equation (1) shows that species *i* impacts log |**Σ**| through its variance 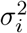 (i.e., the sensitivity of species *i* to perturbations (31)), which expands the volume of perturbed abundances along a given direction (Fig. 2a). In the absence of interspecific effects and correlations in initial perturbations (i.e., **J** and **Σ**_*t*_ are diagonal matrices), **P** = **I** and |**P**| = 1, implying that log |**Σ**| is completely determined by 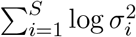. However, in the presence of interspecific effects, 0 ≤ |**P**| ≤ 1 and species correlations impact community sensitivity by decreasing the volume of perturbed abundances (Fig. 2a). That is, given a contribution of individual species 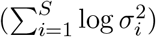, the contribution of species correlations (log |**P**|) can only decrease community sensitivity (i.e., decrease log |**Σ**|) (Fig. 2a). This result suggests that an intuitive way to understand the relative importance of species correlations is to consider the fraction of community sensitivity explained by the contribution of individual species: 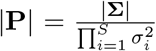. Thus, |**P**| → 1 indicates that community sensitivity is completely explained by the contribution of individual species, whereas |**P**| → 0 indicates that community sensitivity is completely explained by the contribution of species correlations.

To illustrate how we can understand log |**Σ**| in light of 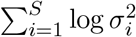 and log |**P**| (Fig. 2b), we use the predator-prey model from equation (2) (*Materials and Methods*). As already shown in Figure 1, depending on the location of the community along this predator-prey cycle, species correlations will be weaker (Figs. 1a, c) or stronger (Figs. 1b, d). Figure 2b shows that **Σ** (black ellipses) indeed captures the changes in the distribution of perturbed abundances at time *t* = *t*_1_ (absence of species correlations) and time *t* = *t*_2_ (presence of species correlations). Importantly, this figure shows that depending on the community state (**N**), log |**Σ**| (purple line) can be dominated by 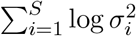 (orange line, *t*_1_, |**P**| ≈ 1) or by log |**P**| (green line, *t*_2_, |**P**| ≪ 1; Fig. 2b).

### Understanding the contribution of species correlated responses using models

We now illustrate how our framework can provide important insights about the state-dependent impact of species correlations (log |**P**|) on community sensitivity to perturbations (log |**Σ**|). To do so, we use three population dynamics models that generate non-equilibrium attractors: a 2-species predator-prey model (equation [2]), a 3-species food chain model (equation [3]), and a 4-species competition model (equation [4]) (*Materials and Methods*). We generate synthetic time series with 250 points ({**N**(*t*)}, *t* = 1, …, 250) from each model and calculate the analytical Jacobian matrix evaluated at each **N**(t) (**J**; *Materials and Methods*). Then, for each point in time, we compute the covariance matrix **Σ** and, finally, the contribution of species correlations as: 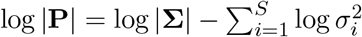.

Figure 3 shows the contribution of species correlations (log |**P**|, green line) through time for each population dynamics model (panel a: 2-species predator-prey model, equation [2]; panel b: 3-species food chain model, equation [3]; panel c: 4-species competition model, equation [4]). Overall, Figure 3 provides three main insights about the impact of species correlations on community sensitivity. First, for all models, we find a large variation in log |**P**| over time. That is, there are points in time when species correlations have no impact on community sensitivity (i.e., |**P**| ≈ 1, for example at time *t*_1_) and points in time when they dominate community sensitivity (i.e., |**P**| ≪ 1, for example at time *t*_4_). Second, we find that the correlation matrix **P** (right panels in Fig. 3) contains information that cannot be directly extracted from the Jacobian matrix **J** (left panels in Fig. 3). Because we convert **J** into **Σ** (and therefore into **P**) by computing a matrix exponential (i.e., *e*^*k***J**^), **P** is the result of a sum of all indirect effects between species. Thus, the sign and strength of species correlations (*ρ*_*ij*_; for example right panel in Fig. 3c) cannot be deduced from the sign and strength of interspecific effects (*j*_*ij*_; for example left panel in Fig. 3c). Finally, we find that, although log |**P**| becomes a complicated nonlinear function of *ρ*_*ij*_ (*i, j* = 1, …, *S*) as *S* becomes large, it is mainly driven by the strongest *ρ*_*ij*_ value in **P** (e.g., right panel in Fig. 3b and Fig. S3). That is, higher values of 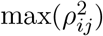 lead to more negative values of log |**P**| (i.e., stronger contribution of species correlations; Fig. S3). We confirm this relationship between 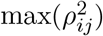 and log |**P**| by randomly generating multiple correlation matrices with up to *S* = 10 species (Fig. S4).

**Figure 3.**
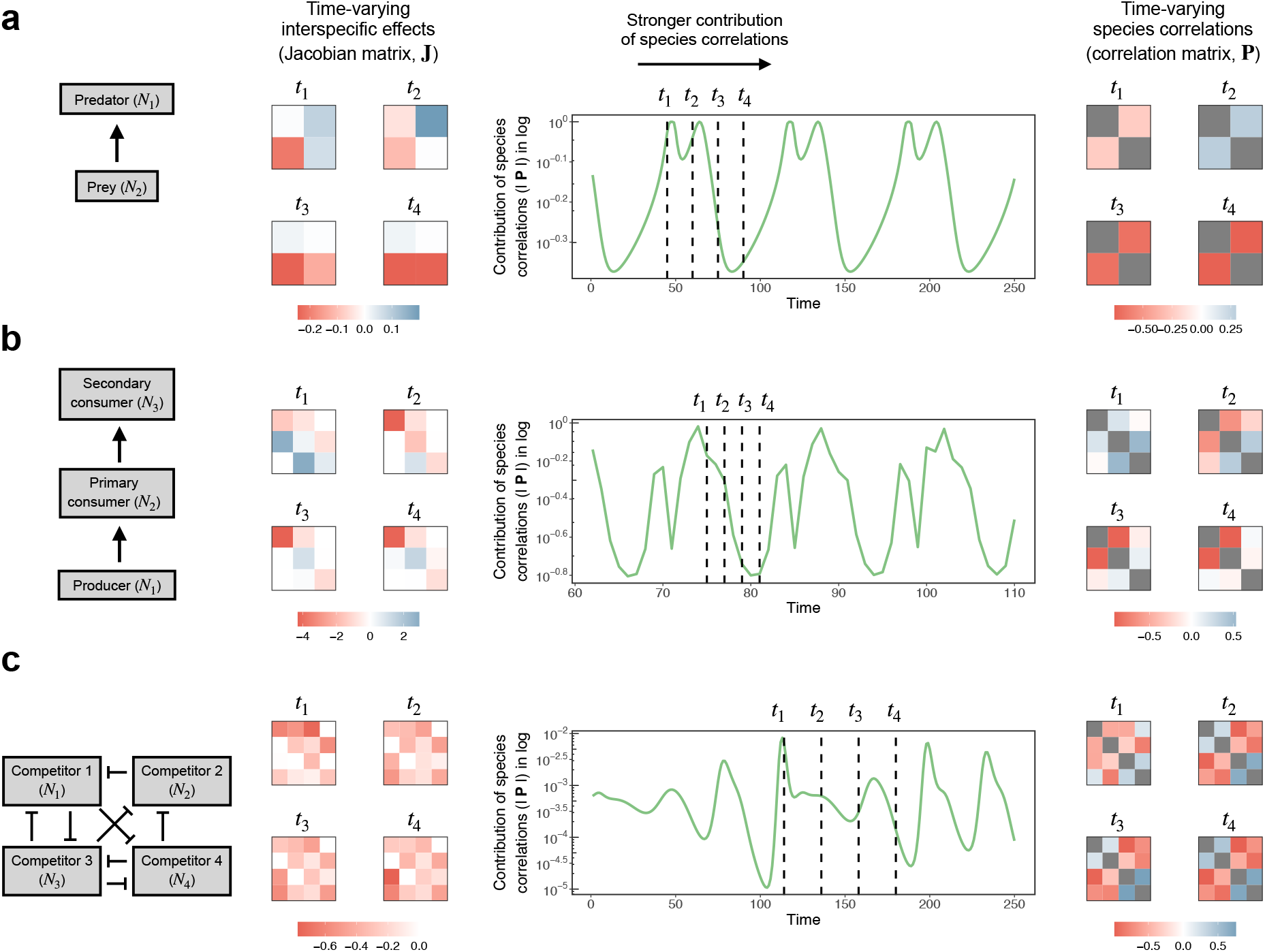
Impact of species correlated responses on community sensitivity changes over time under three population dynamics models. **a**-**c** (center), Contribution of species correlations to community sensitivity (log |**P**|, green line) over time for a 2-species predator-prey (**a**, equation [2]), a 3-species food chain (**b**, equation [3]), and a 4-species competition (**c**, equation [4]) model. Times *t*_1_ through *t*_4_ (vertical dashed lines) depict four arbitrary community states with an increasing contribution of species correlations (i.e., progressively more negative log |**P**|). **a**-**c** (left), Jacobian matrix (**J**) containing time-varying interspecific effects at each of the four states. **a**-**c** (right), Correlation matrix (**P**) containing time-varying species correlated responses to perturbations at each of the four states. Although one or more correlation values in **P** become stronger, there is no clear pattern in how **J** changes from time *t*_1_ to *t*_4_. Note that all diagonal elements in **P** are equal to 1 and are colored in gray.

### Inferring the contribution of species correlated responses from time series

When dealing with empirical communities, there is typically a large uncertainty regarding model structure and parameters (29, 41, 42). Thus, calculating the time-varying Jacobian matrix (**J**) using a parameterized population dynamics model to investigate the impact of species correlations on community sensitivity is rarely feasible. In light of these limitations, we now show how to apply our framework by inferring **J** directly from noisy time-series data without using model equations.

Given a multivariate abundance time series of *S* species, we can infer **J** at each point using a locally weighted state-space regression method known as the S-map (*Materials and Methods*) (21, 43–45). To verify the accuracy of inferring the contribution of species correlations (log |**P**|) from time-series data, we perform the S-map to infer **J** at each point in time for each of the synthetic time series generated from the three population dynamics models (*Materials and Methods*). We then compute the contribution of species correlations at each point in time as: 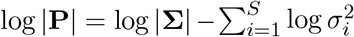. Because empirical time series are usually contaminated with observational noise (43, 44), we apply Gaussian noise to each synthetic time series before performing the S-map (*Materials and Methods*). Importantly, note that our framework focuses on inferring only two quantities: the trace of **J** (i.e., volume expansion rate), which is equivalent to log |**Σ**|, and the trace (in log) of **Σ** (i.e., contribution of individual species). By relying on an accurate inference of just these two quantities, our framework minimizes inference errors, especially in communities with a large number of species (29, 30).

Because we can only infer the elements of **J** up to a constant (29) and the value of log |**P**| depends on *S* and *k* (Figs. S4 and S5), here we focus on qualitatively detecting points in time with extreme (low or high) values of log |**P**| (*Materials and Methods*). Importantly, we find the inferred log |**P**| to be very similar over time to the analytical log |**P**| for all three synthetic time series (Fig. 4). In particular, for all synthetic time series, we obtain an accuracy approximately twice as better as a random guess (i.e., 25%): 43.6% for the 2-species predator-prey model (Fig. 4a), 54.8% for the 3-species food chain model (Fig. 4b), and 53.2% for the 4-species competition model (Fig. 4c). Although this accuracy can decrease with stronger observational noise and under uncertainty in **Σ**_*t*_ and *k* (*Materials and Methods*), we still obtain an accuracy higher than the random expectation of 25% for most cases (Figs. S6 and S7).

**Figure 4.**
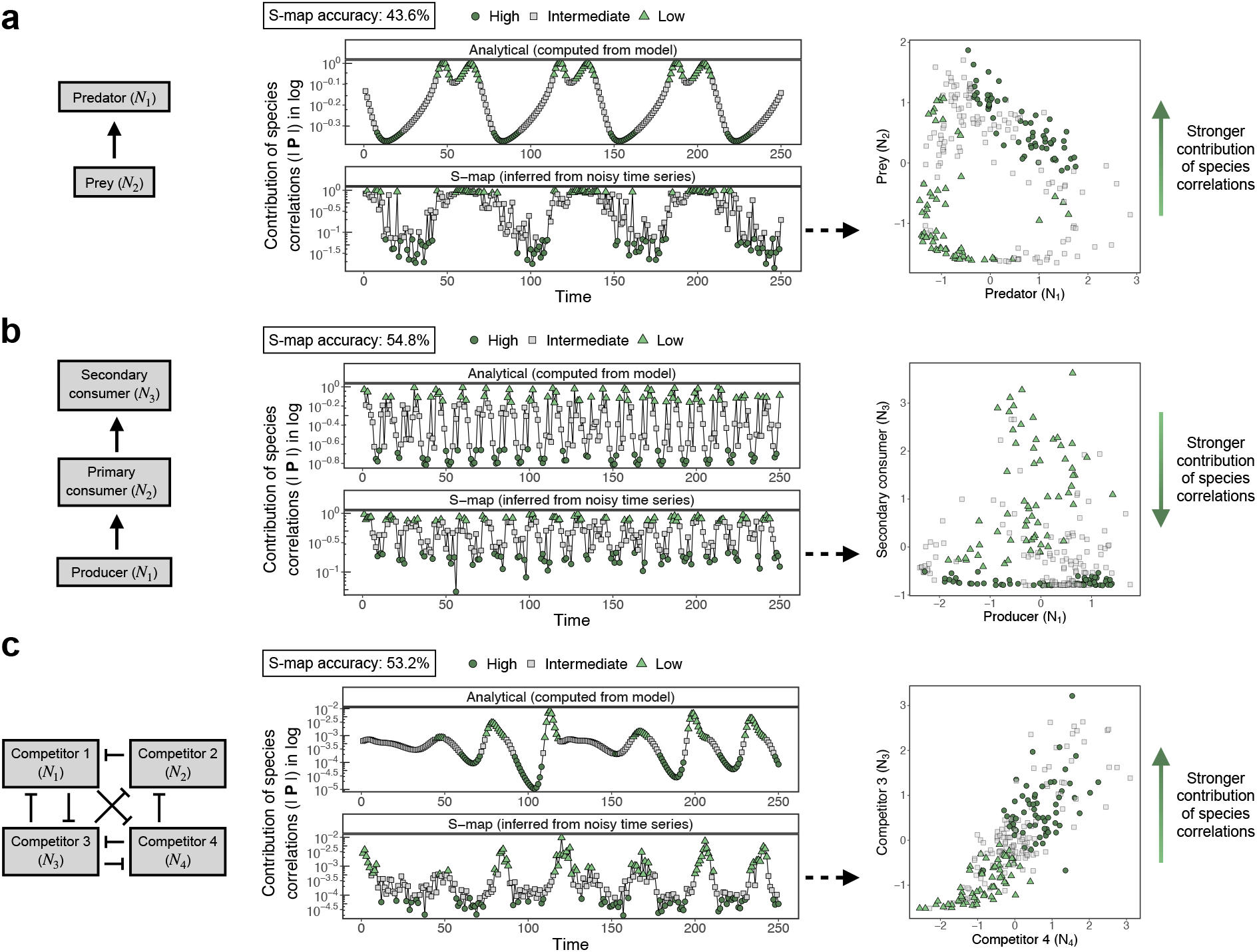
State-dependent impact of species correlated responses can be inferred directly from noisy time series. **a**-**c** (center), For each population dynamics model (**a**, equation [2]; **b**, equation [3]; **c**, equation [4]), the top panel shows the contribution of species correlations (log |**P**|) over time computed analytically from the model equations, whereas the bottom panel shows log |**P**| inferred from the noisy time series with the S-map. Values of log |**P**| for each panel are independently classified as having a high (dark green circles), intermediate (gray squares), or low (light green triangles) contribution to community sensitivity. The S-map accuracy indicates the percentage of high and low points in the bottom panel that match those in the top panel (expected accuracy of random guess: 25%). **a**-**c** (right), State space of species abundances (with noise and normalized) for each model showing that the inferred high and low values of log |**P**| are concentrated along certain community states. For instance, species correlations only reduce community sensitivity when the prey abundance is high under the 2-species model (**a**). Note that for the 3- and 4-species models (**b**-**c**), we show the two species that most clearly separate high and low values of log |**P**|.

Most importantly, we find low and high values of log |**P**| to be concentrated along certain regions of each non-equilibrium attractor (Figs. 4 and S8). For example, for the 2-species predator-prey model, we find that species correlations can only decrease community sensitivity when the prey abundance (*N*_2_) is high (Fig. 4a). We confirm this result with an additional 2-species predator-prey model (Fig. S8) and find similar simple state-dependencies in the 3-species food chain model (Fig. 4b) and in the 4-species competition model (Fig. 4c). These results suggest that the impact of species correlations should only be observed under certain species abundance states, which we can accurately detect using a time-series based inference approach.

### Application to experimental communities

Lastly, we showcase our framework using two empirical communities to gain further insights into how and when species correlations may reduce community sensitivity to perturbations. We use two experimental time series with 2 and 3 species, respectively, that have been shown to exhibit non-equilibrium dynamics under laboratory conditions (25, 26) (*Materials and Methods*). Following our analyses with synthetic time series, we apply the S-map to both time series to infer the contribution of species correlations to community sensitivity over time as: 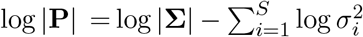. Given our accuracy in detecting regions with extreme (low or high) values of log |**P**| in synthetic time series (Fig. 4), we follow the same procedure for these experimental time series.

We find that the fraction of community sensitivity explained by the contribution of individual species 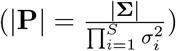 changes considerably across time for both the 2-species (range of |**P**|: 0.77-0.99, Fig. 5a) and the 3-species community (range of |**P**|: 0.64-0.93, Fig. 5b). Note that these fractions depend on the time step *k* (*Materials and Methods*) and can decrease (i.e., contribution of species correlations can increase) when this time step is larger (Figs. S9). This suggests that either the contribution of individual species or of species correlations can dominate the sensitivity of these communities (|**P**| close to 1 or close to 0, respectively) depending on the point in time. In fact, as expected from our analyses with synthetic time series, we find that time series points with low or high log |**P**| (dark or light green points in Figs. 5a, b) are clearly separated in the state space of species abundances. For the 2-species community, we find that prey abundance is higher when the contribution of species correlations is high than when it is low (two sample t-test: *t*(118.7) = 10.02, *p <* 0.0001). This shows a strong resemblance in the separation of low and high log |**P**| between the 2-species experimental community (Fig. 5a) and the 2-species predator-prey models (Figs. 4a and S8). As for the 3-species community, we find that the abundance of the preferred prey is lower when the contribution of species correlations is high than when it is low (two sample t-test: *t*(9.69) = −3.38, *p* = 0.007), confirming again that the impact of species correlations on community sensitivity depends on the abundance of particular species.

**Figure 5.**
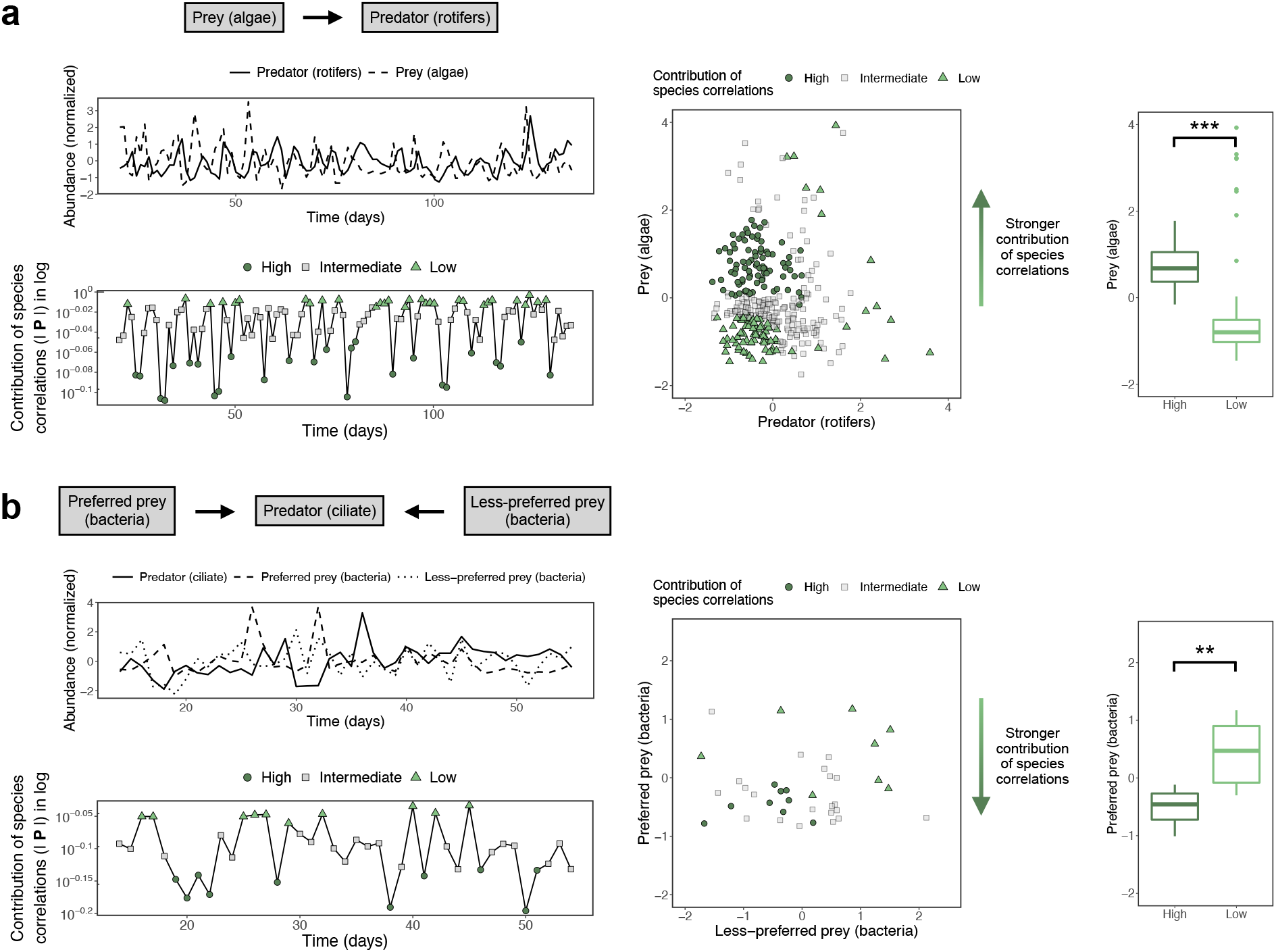
Impact of species correlated responses depends on prey abundance for two experimental communities. **a** (left), The top panel shows a section of the (normalized) abundance time series of a predator (rotifers, solid line) and its prey (algae, dashed line). The bottom panel shows the contribution of species correlations to community sensitivity (log |**P**|) for the same time series section with points being classified as having a high (dark green circles), intermediate (gray squares), or low (light green triangles) contribution. **a** (right), The left panel shows the state space of species abundances for the complete time series with each point colored according to the contribution of species correlations. The right panel shows that the prey abundance (boxplots) is higher for points with a high than with a low contribution (asterisks denote *p <* 0.0001 for a two sample t-test). **b** (left), The top panel shows the (normalized) abundance time series of a predator (ciliate, solid line), its preferred prey (bacteria, dashed line), and its less-preferred prey (bacteria, dotted line). The bottom panel shows log |**P**| over time with points being classified as in **a. b** (right), The left panel shows the state space of prey abundances with each point colored according to the contribution of species correlations. The right panel shows that the preferred prey abundance (boxplots) is lower for points with a high than with a low contribution (asterisks denote *p* = 0.007 for a two sample t-test).

## Discussion

The response of an ecological community to perturbations (e.g., recovery, constancy, sensitivity) is a property that emerges from the responses of its constituent species and the interactions among them (7–10). Thus, understanding how responses at the species-level scale up to shape the response of the whole community has remained an important problem in ecology (6, 7, 46, 47). Here, we tackled this problem when community dynamics are out of equilibrium and, therefore, interspecific effects (i.e., elements of the Jacobian matrix) change over time and have time-varying impacts on whole-community response to perturbations. Our work provides three main insights about how species correlated responses to perturbations impact community response under non-equilibrium dynamics.

First, by developing a framework to decompose the community sensitivity to perturbations into contributions of individual species and of species correlations, we demonstrate that correlations (positive or negative) can only dampen community sensitivity (Fig. 2). Previous work on compensatory dynamics has established that negative correlations increase constancy of total abundance (a measure of whole-community response), whereas positive correlations decrease it (9, 14–16). Although constancy is an important measure of community response, it is most relevant when abundances are assumed to be at equilibrium (16). Under the abundance fluctuations generated by species interactions investigated here, the volume expansion rate of perturbed abundances (29, 30, 39) provides a more meaningful measure of the short-term community response to perturbations. In fact, given that natural communities are constantly perturbed, the short-term species correlations that we explore here can be a potential explanation for the compensatory or synchronous dynamics found at short timescales in several communities (48–50). Overall, based on our framework, there are two ways for perturbed abundances to have a small volume expansion rate in state space: the contribution of individual species is low (i.e., variances in **Σ** are small) or the contribution of species correlations is high (i.e., |**P**| is close to zero). Note that, if variances in **Σ** are large but |**P**| is close to zero, perturbed abundances will stretch along a given direction even though the volume expansion rate is small. Therefore, to understand how perturbations will affect an entire community under non-equilibrium dynamics (29) we need to combine information on the response of individual species (31) with species correlated responses.

Second, we find that the impact of species correlations on community sensitivity changes over time and essentially depends on the maximum squared correlation value in the correlation matrix **P** (Fig. 3). Thus, one strong correlation value in **P** (positive or negative) is generally sufficient to bring |**P**| close to zero, greatly reducing the volume expansion rate (i.e., reducing community sensitivity). This single strong correlation provides a simple mechanism for how a pair of species responding together to perturbations may dampen community sensitivity, although this mechanism may not be enough in large communities (Fig. S4). In addition, we show that the covariance matrix **Σ** results from a sum of all indirect effects between species (equation [6]), suggesting a non-trivial relationship between interspecific effects and species correlated responses. Given the importance of indirect effects for ecological dynamics in species-rich communities (51–53), investigating the links between time-varying interspecific effects and species correlations can be an interesting future direction. Specifically, combining our framework with recent improvements of the S-map approach to infer the time-varying Jacobian matrix of large communities from time-series data can be a fruitful research avenue (44, 45).

Finally, we find that the contribution of species correlations to community sensitivity depends on community state under both models (Fig. 4) and experimental data (Fig. 5). That is, even though community dynamics may follow a non-equilibrium attractor (i.e., perturbed abundances eventually return to the attractor), short-term responses to perturbations may vary considerably along the attractor. In particular, points in time with a high or low contribution of species correlations are not scattered along a non-equilibrium attractor, but rather confined to specific regions of the state space of species abundances. Therefore, our results indicate that species correlated responses may only contribute to decrease community sensitivity under certain values of species abundances (e.g., when prey abundance is high, Figs. 4a and 5a). Previous food web studies have found that species responses after predator removal or introduction depends on the sign and strength of species interactions (17–19). We find an extension of these results for non-equilibrium communities, where state-dependent interspecific effects create state-dependent species correlations. Given that consumer-resource interactions frequently generate cyclic or chaotic dynamics (21, 25–27, 54), such state-dependent impact of species correlations on community response to perturbations may be widespread in natural communities. If this is the case, our framework can potentially be used to plan species removal or introductions (i.e., intentional perturbations) so as to minimize whole-community changes. All in all, our framework allows us to better understand how and when species interactions can dampen community sensitivity to perturbations when we break the standard ecological assumption of equilibrium dynamics.

## Materials and Methods

### Synthetic time series from population dynamics models

We use three synthetic time series generated from population dynamics models with *S* = 2, *S* = 3, and *S* = 4 species to illustrate the impact of species correlations on community sensitivity to perturbations under non-equilibrium dynamics. The first model depicts the population dynamics of a predator (species 1) and its prey (species 2) (33):

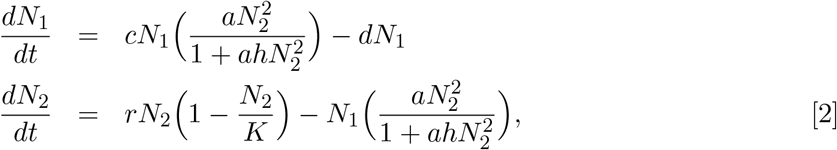

where *c* and *d* are the predator conversion and death rates, *a* is the encounter rate, *h* is the handling time, and *r* and *K* are the prey intrinsic growth rate and carrying capacity. Here, we explore the limit cycle that emerges when *c* = 0.5 *a* = 0.002, *h* = 4, *d* = 0.1 *r* = 0.5, and *K* = 100 (33) (Figs. 1 and S2). The second model depicts a food chain with a producer (species 1), a primary consumer (species 2), and a secondary consumer (species 3) (54, 55):

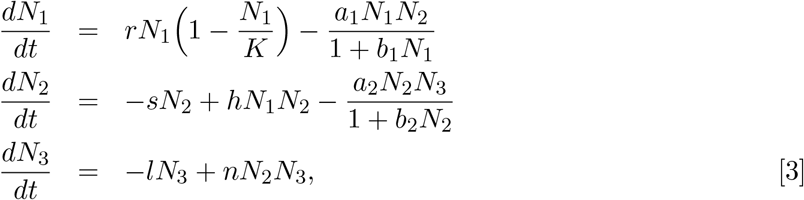

where *r* and *K* are the producer intrinsic growth rate and carrying capacity, *a*_1_ and *a*_2_ are encounter rates, *b*_1_ and *b*_2_ are handling times, *s* and *l* are consumer death rates, and *h* and *n* are consumer conversion rates. We explore the chaotic attractor that arises when *r* = 4.3, *K* = 50, *a*_1_ = 0.1, *b*_1_ = 0.1, *a*_2_ = 0.1, *b*_2_ = 0.1, *s* = 1, *h* = 0.05, *l* = 1, and *n* = 0.03 (55) (Fig. S2). The third model consists of the classic Lotka-Volterra model with competitive interactions (32):

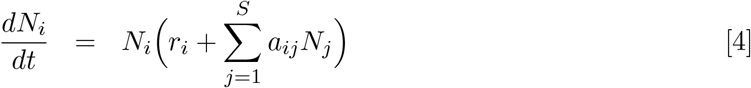

where *r*_*i*_ (*r*_*i*_ *>* 0) is an element of the vector **r** representing the intrinsic growth rate of species *i* and *a*_*ij*_ (*a*_*ij*_ ≤ 0) is an element of the interaction matrix **A** representing the per-capita effect of species *j* on species *i*. We investigate the chaotic attractor that emerges with *S* = 4 and the following parameter values (56) (Fig. S2):

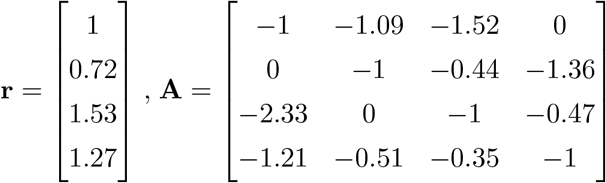

For each model, we first numerically integrate the dynamics using a Runge-Kutta method with a time step of 0.05 and obtain a time series with 5,000 points. Then, we sample equidistant points obtaining a final multivariate time series with 250 points ({**N**(*t*)}, *t* = 1, …, 250). Note that with this protocol we obtain time series that fully sample the attractor of each model (Fig. S2).

### Community sensitivity as volume expansion rate

A generic population dynamics model for a community with *S* species may be written as: 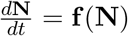, where **N** = [*N*_1_, …, *N*_*S*_]^┬^ is the vector of species abundances and **f** = (*f*_1_, …, *f*_*S*_) (*f*_*i*_: ℝ^*S*^ →ℝ) is a set of functions. At any given state **N**, this community can be affected by a small pulse perturbation **p** = [*p*_1_, …, *p*_*S*_]^┬^ that changes **N** into Ñ (34). The linearized dynamics of such perturbation is given by (*Supporting Information Section 1*) (31, 40):

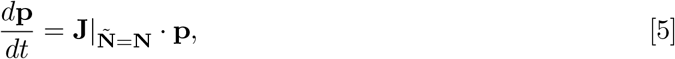

where **J**|_**Ñ** =**N**_ (hereafter **J**) is the Jacobian matrix of partial derivatives with elements 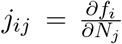 (interspecific effects) evaluated at **N**. Note that **J** is state-dependent as it depends on the community state **N**. Because we typically have no knowledge of how perturbations will affect a given community, we assume that perturbations at an arbitrary time *t* (**p**(*t*)) follow a distribution with mean vector ***µ***_*t*_ and covariance matrix **Σ**_*t*_ (Fig. 2a). Although this is not necessary for our derivation (*Supporting Information Section 1*), we focus on perturbations that are not biased in a given direction (i.e., ***µ***_*t*_ = **0**) and that affect each species equally and independently (i.e., **Σ**_*t*_ = *c***I**, where **I** is the identity matrix and *c* is the perturbation variance). After *k* time steps, the covariance matrix describing the distribution of **p**(*t* + *k*) will be given by (*Supporting Information Section 1*) (31, 35):

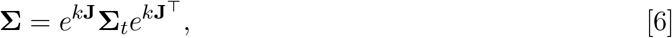

where 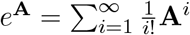 is the exponential of matrix **A**.

Because the determinant of a matrix represents the volume of the *S*-dimensional parallelepiped formed by its row vectors (37, 38), the volume of perturbed abundances at time *t* and *t* + *k* will be proportional to the determinants of the respective covariance matrices: |**Σ**_*t*_| and |**Σ**|. Here, we are interested in the change in volume from time *t* to *t* + *k* and, therefore, consider the volume of initial perturbations to be fixed: |**Σ**_*t*_| = *v*. The determinant of **Σ** is then given by:

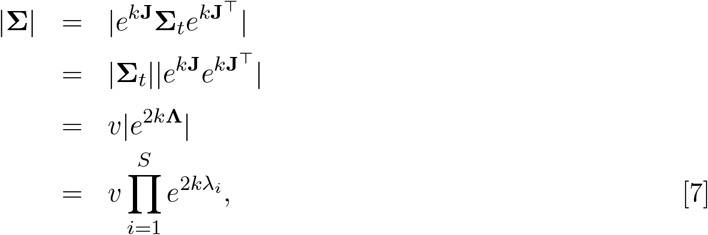

where **Λ** is a diagonal matrix containing the eigenvalues of **J** (*λ*_1_, …, *λ*_*S*_). Taking the logarithm of both sides of equation [7], we obtain a connection to the volume expansion rate:

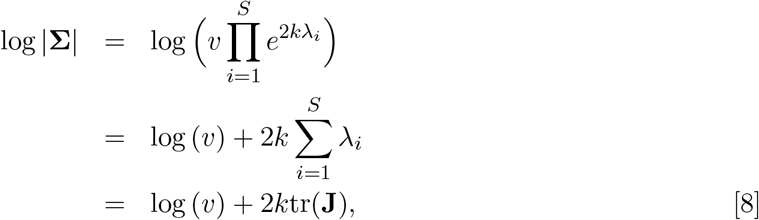

where tr(**J**) is the trace of **J**. Note that 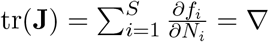. **f** (**N**) represents the divergence of the vector field **f** around **N** (i.e., volume expansion rate) (29, 30, 40). Therefore, log |**Σ**| is equivalent to tr(**J**) and represents the community sensitivity to pulse perturbations on abundances.

### Computing the contribution of species correlated responses from time series

For each synthetic time series generated from population dynamics models, we compute the time-varying contribution of species correlations (log |**P**|) in two different ways. Recall that, as described in equation [1], log |**P**| is given by the difference between the community sensitivity (log |**Σ**|) and the contribution of individual species 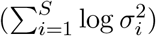. First, we compute log |**P**| analytically by calculating the Jacobian matrix **J** from the model equations (equations [2], [3], and [4]) and evaluating it at each time series point **N**(*t*). Second, we compute log |**P**| by inferring **J** directly from the time series with the S-map method (see next section). To compute **Σ**, we assume minimal knowledge of the distribution and evolution of perturbations and set **Σ**_*t*_ = **I** and *k* to be a fixed value over time (*k* = 3 for 2-species predator-prey and 4-species competition models and *k* = 0.5 for 3-species food chain model). Note that these values of *k* are proportional to the average rate of change (i.e., 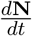) of each model and that using other values of *k* gives qualitatively similar results for all models (Fig. S5). Then, for each point in time, we use **Σ** to calculate the contribution of species correlations as: 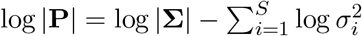.

Instead of focusing on the exact values of log |**P**| over time, here we focus on qualitatively detecting community states with extreme (low or high) log |**P**| values. That is, after computing log |**P**| analytically (hereafter analytical log |**P**|) or with the S-map (hereafter inferred log |**P**|), we classify each time series point as having a low (i.e., log |**P**| higher than its 75th percentile) or a high (i.e., log |**P**| lower than its 25th percentile) contribution of species correlations. We perform this classification independently for the analytical and inferred log |**P**|. Then, to test the accuracy of inferring log |**P**| with the S-map, for each of the three synthetic time series, we compute the percentage of points classified as having a low (or high) analytical log |**P**| that also have a low (or high) inferred log |**P**|. Note that if we were randomly classifying points as having a low or high log |**P**| (instead of using the inferred log |**P**|), our accuracy of matching the analytical classification would be, on average, 25%.

### Inferring the time-varying Jacobian matrix with the S-map

Here we describe the S-map, a locally weighted state-space regression method that has been shown to provide accurate inferences of the time-varying Jacobian matrix (**J**) from abundance time series (21, 43–45). We perform the S-map using the rEDM package in R to sequentially infer **J** using only past time-series data. Given a multivariate time series containing the abundances of *S* species over *T* time points ({**N**(*t*)}, *t* = 1, …, *T*), the S-map method allows us to infer **J** at each point. This method is based on fitting a linear regression of the following form to the time series: 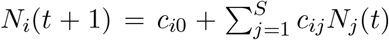, where 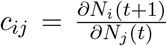 is a discrete-time approximation of the Jacobian matrix element *j*_*ij*_. However, fitting this linear regression would not capture the state-dependent nature of **J**, that is, **J** fundamentally depends on **N** under non-equilibrium dynamics (e.g., left panels in Fig. 3). Thus, the S-map consists of fitting this linear regression locally for each target point **N**(*t*^∗^) by giving a stronger weight to points that are closer to it in state space. This is done by finding a solution for **c** in **b** = **Ac**, where *b*_*t*_ = *w*_*t*_*N*_*i*_(*t* + 1), 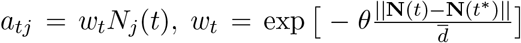, and 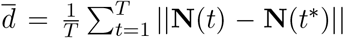. Thus, b ∈ ℝ^*T*^ contains the abundances at *t* + 1 weighted by the relative distance of each point to the target point, **A** ∈ ℝ^*T* ×(*S*+1)^ is the weighted data matrix of abundances at *t*, and **c** ∈ ℝ^*S*+1^ estimates the *i*th row of **J** at **N**(*t*^∗^) as well as an intercept term. We obtain the solution for **c** via singular value decomposition (21), which is equivalent to the ordinary least squares solution (44). Finally, the parameter *θ* tunes how strongly the regression is localized around each target point **N**(*t*^∗^) and we select its value via abundance predictions with leave-one-out cross-validation (44). That is, for a given *θ* value, we fit the S-map to the time series after removing one of its points (e.g., **N**(*t*′)), use **J** at **N**(*t*′ − 1) to predict **N**(*t*′), and repeat this procedure by removing each of the *T* points. We then select the *θ* value that minimizes the mean prediction error across all *T* abundance predictions.

We perform the S-map on the three synthetic time series to test the accuracy of this method in inferring the time-varying contribution of species correlations (log |**P**|). To do so, we first apply Gaussian noise to each synthetic time series. Specifically, for each species *i* and time *t*, we transform *N*_*i*_(*t*) into *N*_*i*_(*t*) + *𝒩* (*µ* = 0, *σ*^2^ = [*δN*_*i*_(*t*)]^2^), where *δ* = 0.1 (see Fig. S6 for *δ* = 0.2). Then, for each time series, we fit the S-map with the leave-one-out cross-validation procedure to select the best *θ* parameter and use this value of *θ* to fit the S-map to the whole time series and obtain **J** at each point in time. Note that, because species abundances typically vary in scale, we normalize each noisy time series to zero mean and unit standard deviation prior to performing the S-map (20, 21, 44). Following the analysis with the analytical matrix **J** (see previous section), we use each inferred matrix **J** to compute **Σ** by setting **Σ**_*t*_ = **I** and *k* to be a fixed value over time. We also perform these analyses by adding noise to **Σ**_*t*_ and *k* to test the robustness of our framework to uncertainty in these quantities (Fig. S7).

### Inferring the contribution of species correlated responses in experimental communities

Here we describe how we apply our framework to two experimental non-equilibrium time series. The first time series represents a 2-species microcosm community containing the rotifer *Brachionus calyciflorus* as a predator species and the alga *Monoraphidium minutum* as its prey species (26). This community has been shown to exhibit population cycles under constant experimental conditions for approximately 300 predator generations (26). Due to missing data and unequal sampling intervals, we interpolate the original time series using cubic hermite interpolation to obtain a final time series with equidistant time intervals of 1.045 day. The second time series represents a 3-species chemostat community containing the ciliate *Tetrahymena pyriformis* as a predator species and the bacteria *Pedobacter* sp. and *Brevundimonas* sp. as its two prey species (25). It has been demonstrated that under some experimental conditions, this community can exhibit chaotic dynamics (i.e., positive Lyapunov exponent) (25). Because this time series contains equal sampling intervals of 1 day, we did not use an interpolation procedure for it. We use the time series starting from day 14 due to several missing data points prior to this day. The final time series length was of 358 points for the 2-species community and 42 points for the 3-species community. Note that 42 points is more than enough to apply the S-map to a 3-species community (57).

After the time series treatment described above, we apply the S-map (see previous section) to both time series to infer log |**P**| over time. Because of scale differences in species abundances, we normalize each time series to zero mean and unit standard deviation before performing the S-map. Following our analyses with synthetic time series, we select *θ* with leave-one-out cross-validation and use the selected value to fit the S-map to the whole time series and compute 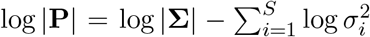 at each point in time. Note that we use each inferred matrix **J** to compute **Σ** by setting **Σ**_*t*_ = **I** and *k* = 1 as we have no information about how perturbations impact these communities (see Fig. S9 for *k* = 3). Finally, we classify each time series point as showing a low (i.e., log |**P**| higher than its 75th percentile) or a high (i.e., log |**P**| lower than its 25th percentile) contribution of species correlations.

## Acknowledgments

We thank C. Long, J. Deng, V. Dakos, G. Sugihara, T. Lieberman, and D. Rothman for insightful discussions and suggestions regarding this work. L.P.M. was supported by MIT ESI and the Martin Family Society of Fellows for Sustainability. S.S. was supported by NSF DEB-2024349 and the MIT Sea Grant Program.

