## Supporting Information for "Understanding the state-dependent impact of species correlated responses on community sensitivity to perturbations"

### 1 Derivation of expectation and covariance matrix of perturbed abundances

Here, we derive the linear dynamics of small perturbations as well as the expectation and covariance matrix describing the distribution of these perturbations under non-equilibrium dynamics. Following the main text, we write the generic population dynamics of a community with $S$ species as: $\frac{d\mathbf{N}}{dt} = \mathbf{f}(\mathbf{N})$, where $\mathbf{N} = [N_1, ..., N_S]^\top$ is the vector of species abundances and $\mathbf{f} = (f_1, ..., f_S)$ ($f_i$: $\mathbb{R}^S \to \mathbb{R}$) is a set of functions describing species abundance growth rates. Note that each $f_i$ also depends on a set of parameters, which we consider to be fixed over time. At any given state $\mathbf{N}$, a pulse perturbation $\mathbf{p} = [p_1, ..., p_S]^\top$ may change species abundances from $\mathbf{N}$ into $\tilde{\mathbf{N}}$ (i.e., $\tilde{\mathbf{N}} = \mathbf{N} + \mathbf{p}$) (1). We can obtain the linearized dynamics of a small perturbation $\mathbf{p}$ by computing the Taylor expansion of $\frac{d\tilde{\mathbf{N}}}{dt}$ around $\mathbf{N}$ (2, 3):

$$\frac{d\tilde{\mathbf{N}}}{dt} = \mathbf{f}(\mathbf{N}) + \frac{\partial \mathbf{f}}{\partial \tilde{\mathbf{N}}}\Big|_{\tilde{\mathbf{N}}=\mathbf{N}} \cdot (\tilde{\mathbf{N}} - \mathbf{N}) + O(\mathbf{p}^\top \mathbf{p}), \quad \text{[S1]}$$

where $\frac{\partial \mathbf{f}}{\partial \mathbf{N}} = \mathbf{J}$ is the Jacobian matrix of partial derivatives with $j_{ij} = \frac{\partial f_i}{\partial N_j}$ (interspecific effects), which is evaluated at $\mathbf{N}$. If $\mathbf{p}$ is small, we can approximate its dynamics by taking just the linear term (i.e., ignoring higher-order terms):

$$\begin{aligned} \frac{d\tilde{\mathbf{N}}}{dt} &= \mathbf{f}(\mathbf{N}) + \frac{\partial \mathbf{f}}{\partial \tilde{\mathbf{N}}}\Big|_{\tilde{\mathbf{N}}=\mathbf{N}} \cdot (\tilde{\mathbf{N}} - \mathbf{N}) \\ \frac{d\mathbf{N}}{dt} + \frac{d\mathbf{p}}{dt} &= \frac{d\mathbf{N}}{dt} + \mathbf{J}|_{\tilde{\mathbf{N}}=\mathbf{N}} \cdot \mathbf{p} \\ \frac{d\mathbf{p}}{dt} &= \mathbf{J}|_{\tilde{\mathbf{N}}=\mathbf{N}} \cdot \mathbf{p}. \end{aligned} \quad \text{[S2]}$$

Therefore, the dynamics of a small perturbation $\mathbf{p}$ can be approximated by the linear equation above (2, 3). Note that we have not assumed the existence of an equilibrium here (i.e., $\mathbf{N} = \mathbf{N}^*$ with $\mathbf{f}(\mathbf{N}^*) = \mathbf{0}$) and, therefore, equation [S2] is valid even when abundances are changing over time (e.g., under non-equilibrium dynamics). Importantly, also note that $\mathbf{J}$ can change over time because it depends on species abundances ($\mathbf{N}$; i.e., state-dependent). For a given $\mathbf{N}$ vector at a given time $t$, we can obtain the solution for equation [S2] over a short time period $k$ by assuming that $\mathbf{J}$ does not change much from $t$ to $t + k$: $\mathbf{p}(t + k) = e^{k\mathbf{J}}\mathbf{p}(t)$, where $e^{\mathbf{A}} = \sum_{i=1}^{\infty} \frac{1}{i!}\mathbf{A}^i$ is the exponential of matrix $\mathbf{A}$ (3, 4).

As described in the main text, we often have no knowledge of how perturbations will affect a given community at a given time. Thus, we assume that pulse perturbations at an arbitrary time $t$ ($\mathbf{p}(t)$) follow a distribution with mean vector $\boldsymbol{\mu}_t$ and covariance matrix $\boldsymbol{\Sigma}_t$ (Fig. 2a). Although this is not necessary for the derivations below, we focus on perturbations that are not biased in a given direction (i.e., $\boldsymbol{\mu}_t = \mathbf{0}$) and perturbations that affect each species equally and independently (i.e., $\boldsymbol{\Sigma}_t = c\mathbf{I}$, where $c$ is a constant and $\mathbf{I}$ is the identity matrix). By defining $\mathbf{M} = e^{k\mathbf{J}}$, we can derive the mean vector of the distribution of perturbations at time $t+k$ (3):

$$\begin{aligned} \mathbb{E}[\mathbf{p}(t+k)] &= \mathbb{E}[\mathbf{M}\mathbf{p}(t)] \\ &= \mathbf{M}\mathbb{E}[\mathbf{p}(t)] \\ &= \mathbf{M}\boldsymbol{\mu}_t. \end{aligned} \tag{S3}$$

Note that if $\boldsymbol{\mu}_t = \mathbf{0}$, then $\mathbb{E}[\mathbf{p}(t+k)] = \mathbf{0}$ (i.e., perturbations remain unbiased). In the special case where $\mathbf{p}(t)$ follows a multivariate normal distribution, $\mathbf{p}(t+k)$ also follows a multivariate normal distribution because $\mathbf{M}\mathbf{p}(t)$ is a weighted sum of normal distributions. Most importantly, we can also derive the covariance matrix of $\mathbf{p}(t+k)$, which we denote as $\boldsymbol{\Sigma}$ (3, 4):

$$\begin{aligned} \boldsymbol{\Sigma} &= \mathbb{E}[(\mathbf{p}(t+k) - \mathbb{E}[\mathbf{p}(t+k)])(\mathbf{p}(t+k) - \mathbb{E}[\mathbf{p}(t+k)])^\top] \\ &= \mathbf{M}\mathbb{E}[(\mathbf{p}(t) - \mathbb{E}[\mathbf{p}(t)])(\mathbf{p}(t) - \mathbb{E}[\mathbf{p}(t)])^\top]\mathbf{M}^\top \\ &= \mathbf{M}\boldsymbol{\Sigma}_t\mathbf{M}^\top. \end{aligned} \tag{S4}$$

Thus, $\boldsymbol{\Sigma}$ is obtained via a transformation of the initial covariance matrix ($\boldsymbol{\Sigma}_t$) due to interspecific effects present in the Jacobian matrix $\mathbf{J}$ (Fig. 2a). Note that, in addition to knowing $\mathbf{J}$, knowledge of $\boldsymbol{\Sigma}_t$ and $k$ is required to compute $\boldsymbol{\Sigma}$. The accuracy of $\boldsymbol{\Sigma}$ in describing the distribution of perturbed abundances has been verified under equilibrium dynamics (4) and in the next section we describe our simulations that confirm this accuracy under non-equilibrium dynamics (Fig. S1).

#### 2 Accuracy of covariance matrix in describing perturbed abundances

We perform perturbation simulations to verify the accuracy of the covariance matrix $\boldsymbol{\Sigma} = e^{k\mathbf{J}}\boldsymbol{\Sigma}_t e^{k\mathbf{J}^\top}$ (see previous section) in describing the distribution of perturbed abundances ($\tilde{\mathbf{N}} = \mathbf{N} + \mathbf{p}$). To do so, we use the three synthetic multivariate time series with 250 points ($\{\mathbf{N}(t)\}$, $t = 1, ..., 250$) generated from the population dynamics models described in the main text (see *Materials and Methods* section). For each time series, we apply 300 pulse perturbations ($\mathbf{p} \sim \mathcal{N}(\boldsymbol{\mu}_t, \boldsymbol{\Sigma}_t)$) at each state $\mathbf{N}(t)$. We assume that perturbations are independent for each species (i.e., covariances in $\boldsymbol{\Sigma}_t$ are zero) and are centered in $\mathbf{N}(t)$ (i.e., $\boldsymbol{\mu}_t$ is zero). We also assume that the standard deviation of perturbations (i.e., square root of diagonal elements of $\mathbf{\Sigma}_t$) is the same for every species and is set as 15% of the mean standard deviation of species abundances: $0.15\frac{1}{S}\sum_{i=1}^{S}\sigma_{N_i}$, where $\sigma_{N_i}$ is the standard deviation of $N_i$ for the whole time series. After applying perturbations, we evolve each perturbed state $\tilde{\mathbf{N}}$ over time for $k$ time steps. An example of these perturbed abundances ($\tilde{\mathbf{N}}$) at the initial time $t$ and final time $t+k$ can be seen in Fig. 2b in the main text (initial: light purple points; final: dark purple points).

For each time series and each state $\mathbf{N}(t)$, we compute $\mathbf{\Sigma}$ using the analytical Jacobian matrix $\mathbf{J}$ evaluated at $\mathbf{N}(t)$ as well as $k$ and $\mathbf{\Sigma}_t$ used in the perturbation simulations. Then, we compute the data covariance matrix $\mathbf{S}$ using the 300 perturbed abundances at time $t+k$ ($\tilde{\mathbf{N}}(t+k)$). That is, the $ij$th element of $\mathbf{S}$ contains the covariance between species $i$ ($\tilde{N}_i(t+k)$) and species $j$ ($\tilde{N}_j(t+k)$) computed from the set of perturbed abundances (e.g., dark purple points in Fig. 2b in the main text). Fig S1 shows, for each population dynamics model, the elements of $\mathbf{S}$ (red lines) and of $\mathbf{\Sigma}$ (blue lines) over time. We find a strong correlation over time between the elements of these two matrices for all models: 2-species predator-prey model: 0.87 ± 0.10 (mean ± standard deviation), 3-species food chain model: 0.91 ± 0.04, and 4-species competition model: 0.88 ± 0.03. These strong correlations confirm that $\mathbf{\Sigma}$ captures how the distribution of perturbed abundances changes over time for each state along a non-equilibrium attractor.

### 3 Determinant of covariance matrix

In the main text, we state that the determinant of a covariance matrix can always be written as the product of variances times the determinant of the correlation matrix. Here, we provide a mathematical proof for this statement and also prove that the determinant of the correlation matrix is always between 0 and 1. Let us define a generic $S\times S$ covariance matrix as the following symmetric and positive semi-definite matrix:

$$\mathbf{\Sigma} = \begin{bmatrix} \sigma_1^2 & \sigma_{12}^2 & \cdots & \sigma_{1S}^2 \\ \sigma_{12}^2 & \sigma_2^2 & \cdots & \sigma_{2S}^2 \\ \vdots & \vdots & \ddots & \vdots \\ \sigma_{1S}^2 & \sigma_{2S}^2 & \cdots & \sigma_{SS}^2 \end{bmatrix},$$

where $\sigma_i^2$ is the variance of variable $i$ and $\sigma_{ij}^2$ is the covariance between variables $i$ and $j$. Note that the correlation between $i$ and $j$ is given by: $\rho_{ij} = \frac{\sigma_{ij}^2}{\sigma_i\sigma_j}$. This allows us to rewrite $\mathbf{\Sigma}$ as:

$$\mathbf{\Sigma} = \begin{bmatrix} \sigma_1^2 & \rho_{12}\sigma_1\sigma_2 & \cdots & \rho_{1S}\sigma_1\sigma_S \\ \rho_{12}\sigma_1\sigma_2 & \sigma_2^2 & \cdots & \rho_{2S}\sigma_2\sigma_S \\ \vdots & \vdots & \ddots & \vdots \\ \rho_{1S}\sigma_1\sigma_S & \rho_{2S}\sigma_2\sigma_S & \cdots & \sigma_S^2 \end{bmatrix}.$$

Let us denote the determinant of an arbitrary matrix $\mathbf{A}$ as $|\mathbf{A}|$. Now, we can use the fact that if a constant $c$ multiplies the entire $i$th column (or row) of $\mathbf{A}$, then: $|\mathbf{A}| = c|\mathbf{A}'|$, where $\mathbf{A}'$ is matrix $\mathbf{A}$ after dividing all elements in column (or row) $i$ by $c$. Because each $\sigma_i$ multiplies the entire $i$th column of $\mathbf{\Sigma}$, we have that:

$$|\mathbf{\Sigma}| = \prod_{i=1}^{S} \sigma_i \cdot \begin{vmatrix} \sigma_1 & \rho_{12}\sigma_1 & \cdots & \rho_{1S}\sigma_1 \\ \rho_{12}\sigma_2 & \sigma_2 & \cdots & \rho_{2S}\sigma_2 \\ \vdots & \vdots & \ddots & \vdots \\ \rho_{1S}\sigma_S & \rho_{2S}\sigma_S & \cdots & \sigma_S \end{vmatrix} .$$

In addition, because each $\sigma_i$ multiplies the entire $i$th row of $\mathbf{\Sigma}$, we have that:

$$|\mathbf{\Sigma}| = \prod_{i=1}^{S} \sigma_i^2 \cdot \begin{vmatrix} 1 & \rho_{12} & \cdots & \rho_{1S} \\ \rho_{12} & 1 & \cdots & \rho_{2S} \\ \vdots & \vdots & \ddots & \vdots \\ \rho_{1S} & \rho_{2S} & \cdots & 1 \end{vmatrix} = \prod_{i=1}^{S} \sigma_i^2 \cdot |\mathbf{P}| ,$$

where $\mathbf{P}$ is the correlation matrix. Thus, this proves that $|\mathbf{\Sigma}| = \prod_{i=1}^{S} \sigma_i^2 \cdot |\mathbf{P}|$. In particular, if we take the logarithm of this expression, we obtain equation [1] in the main text, which consists of our decomposition of the community sensitivity to perturbations into contributions of individual species and of species correlations. In addition, we can prove that $0 \leq |\mathbf{P}| \leq 1$ and, therefore, species correlations can only decrease community sensitivity. Without correlations (i.e., $\rho_{ij} = 0$ $\forall\ i \neq j$), we have that $\mathbf{P} = \mathbf{I}$, where $\mathbf{I}$ is the identity matrix. Because in this case $\mathbf{P}$ is a diagonal matrix, its determinant will be given by the product of its diagonal elements: $|\mathbf{P}| = 1$. This gives an upper bound for $|\mathbf{P}|$ as correlations will always decrease this determinant. To obtain the lower bound, we note that because $\mathbf{P}$ is positive semi-definite, its determinant is always nonnegative. Therefore, we have proved that $0 \leq |\mathbf{P}| \leq 1$.

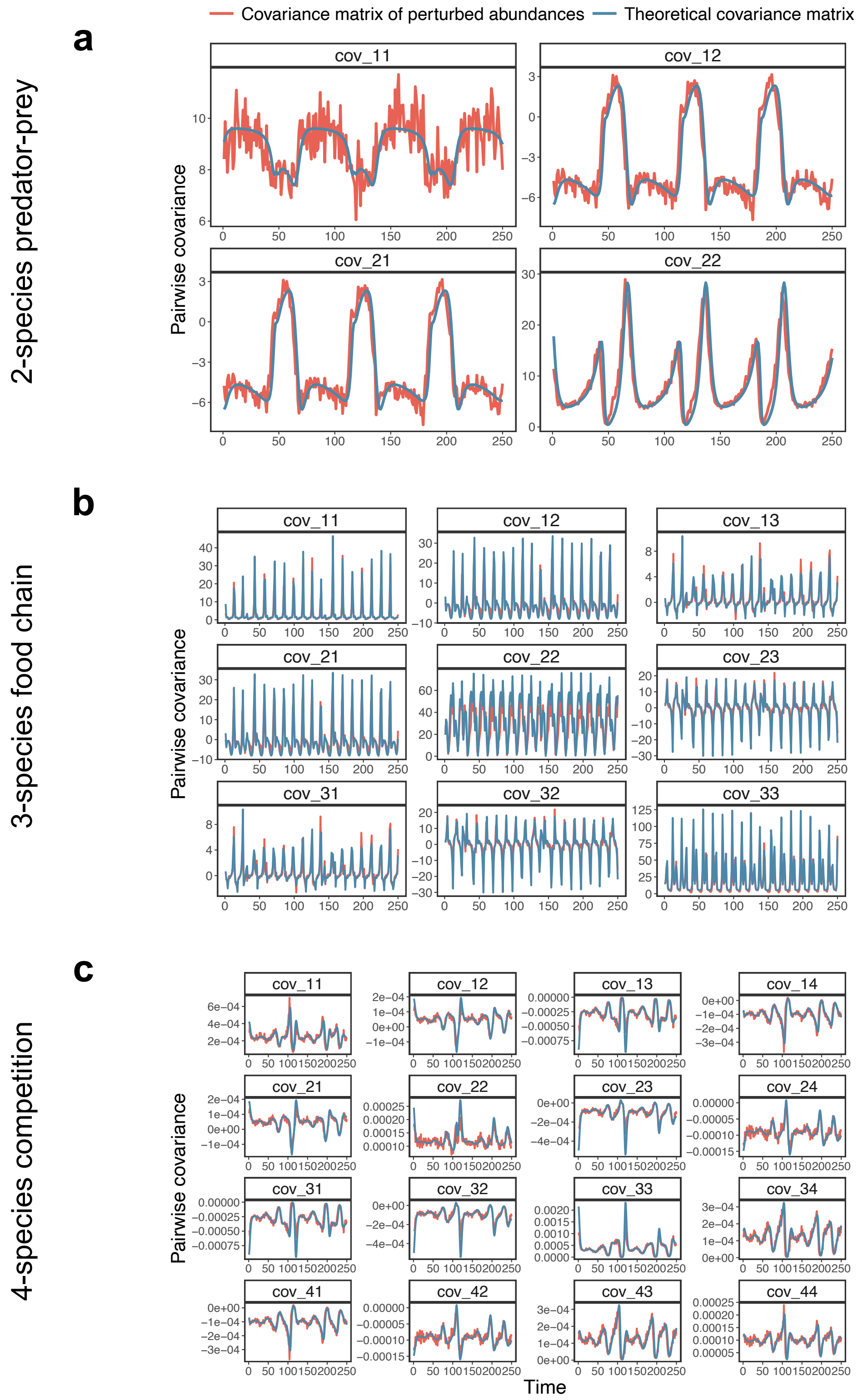


**Figure S1. Accuracy of covariance matrix $\boldsymbol{\Sigma}$ in describing the distribution of perturbed abundances under non-equilibrium dynamics. a-c**, Each panel shows the same element of $\mathbf{S}$ (data covariance matrix, red line) and of $\boldsymbol{\Sigma}$ (blue line) over time. The mean ($\pm$ standard deviation) correlation between an element of $\mathbf{S}$ and an element of $\boldsymbol{\Sigma}$ (correlation between red and blue lines) is: $0.87 \pm 0.10$ for the 2-species predator-prey model (**a**, equation [2] in the main text), $0.91 \pm 0.04$ for the 3-species food chain model (**b**, equation [3] in the main text), and $0.88 \pm 0.03$ for the 4-species competition model (**c**, equation [4] in the main text).

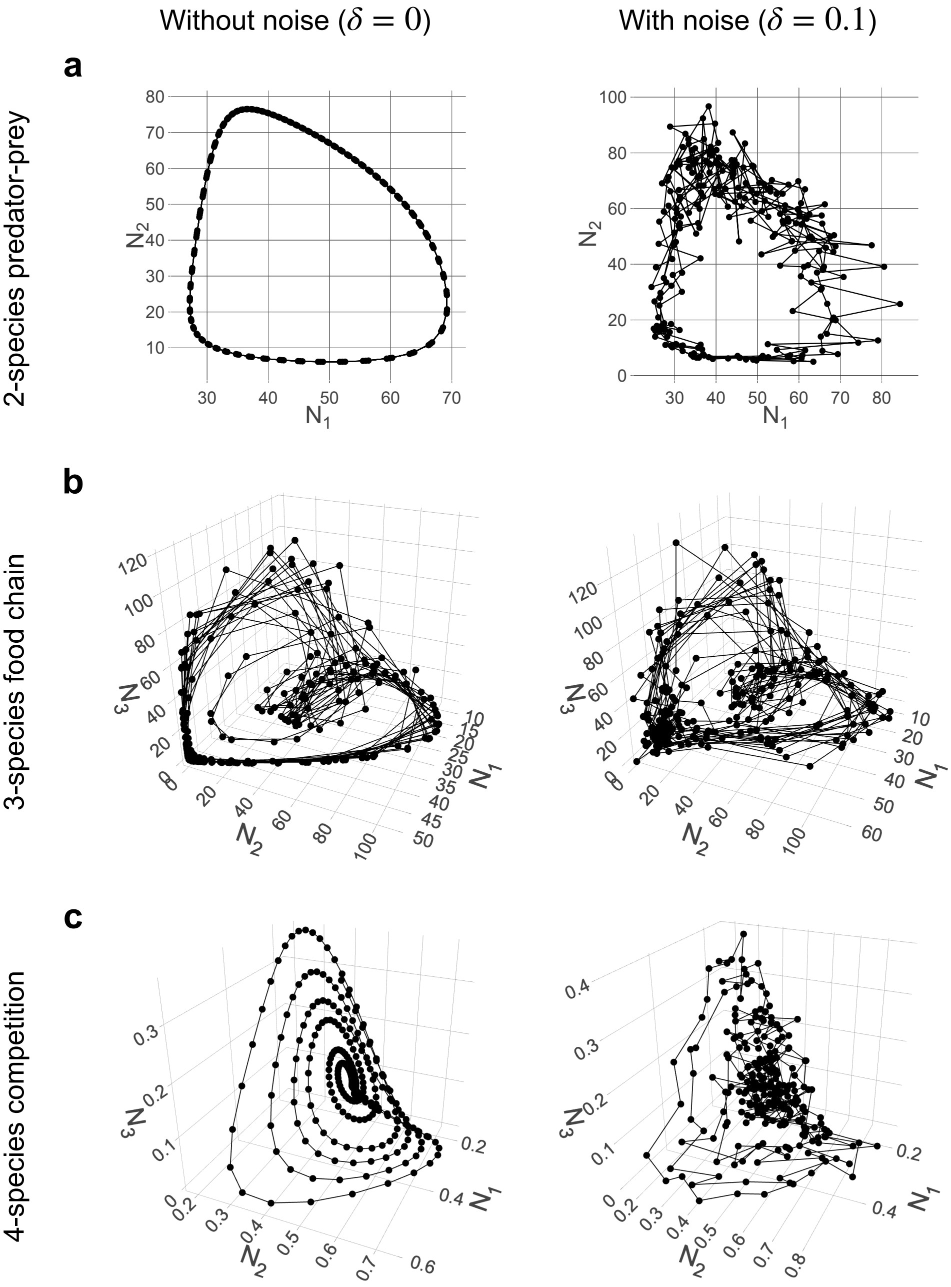


**Figure S2. Non-equilibrium attractors in state space corresponding to each synthetic time series generated by a population dynamics model. a-c**, Each plot shows the 250 points ($\{\mathbf{N}(t)\}$, $t = 1, ..., 250$) generated by numerically integrating the indicated model and then sampling equidistant points. For the left column we did not add noise to the abundances, whereas for the right column we add 10% of Gaussian noise as described in the *Materials and Methods* section in the main text. In our analyses, we compute the analytical Jacobian matrix $\mathbf{J}$ from the attractors without noise (left) and infer $\mathbf{J}$ with the S-map from the attractors with noise (right). Note that we only show the abundances of species 1, 2, and 3 for the 4-species model.

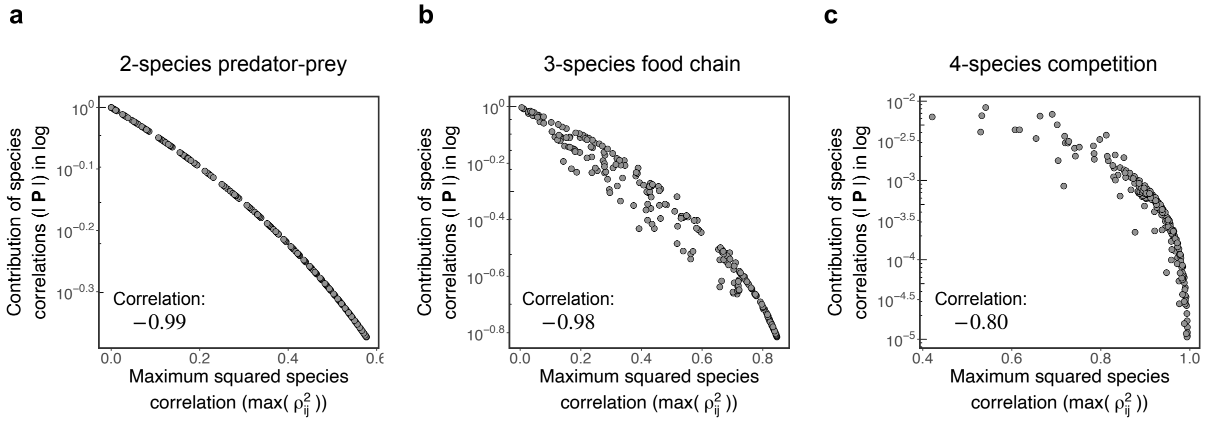


**Figure S3. Contribution of species correlations ($\log|\mathbf{P}|$) is driven by the maximum squared correlation ($\max(\rho_{ij}^2)$). a-c**, Each plot shows $\log|\mathbf{P}|$ and $\max(\rho_{ij}^2)$ computed at each state ($\mathbf{N}(t)$; each point corresponds to a state) of the synthetic time series generated by the indicated population dynamics model. Note that the correlation matrix $\mathbf{P}$ is computed from the covariance matrix $\mathbf{\Sigma}$, which is in turn computed from the analytical Jacobian matrix $\mathbf{J}$ (see *Materials and Methods* in the main text). The correlation between $\log|\mathbf{P}|$ and $\max(\rho_{ij}^2)$ is: $-0.99$ for the 2-species predator-prey model (**a**, equation [2] in the main text), $-0.98$ for the 3-species food chain model (**b**, equation [3] in the main text), and $-0.80$ for the 4-species competition model (**c**, equation [4] in the main text).

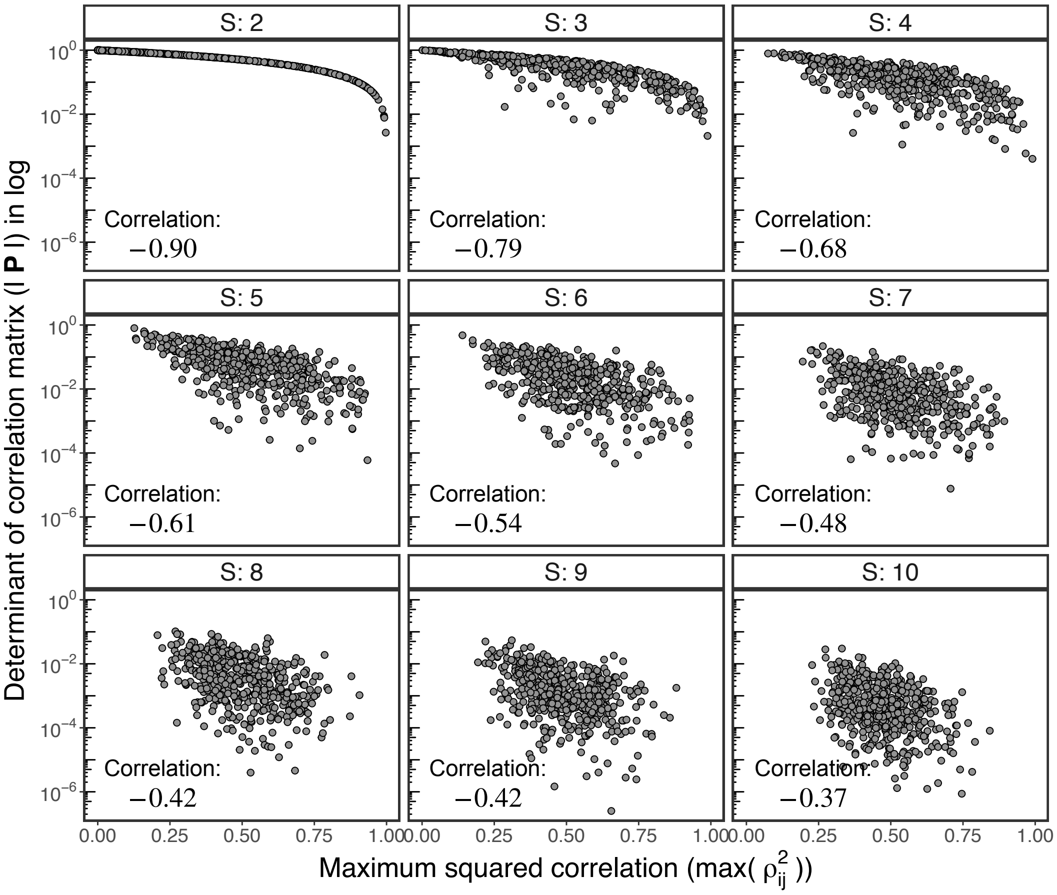


**Figure S4. Log of determinant of correlation matrix ($\log|\mathbf{P}|$) decreases with the maximum squared correlation ($\max(\rho_{ij}^2)$) for random correlation matrices.** Each panel shows $\log|\mathbf{P}|$ as a function of $\max(\rho_{ij}^2)$ for 500 randomly generated correlation matrices (each point denotes one matrix) with a given dimension $S$. Each matrix $\mathbf{P}$ was sampled uniformly over the space of positive definite correlation matrices. Note that, although the correlation between $\log|\mathbf{P}|$ and $\max(\rho_{ij}^2)$ is strong for all values of $S$, it becomes weaker as $S$ increases.

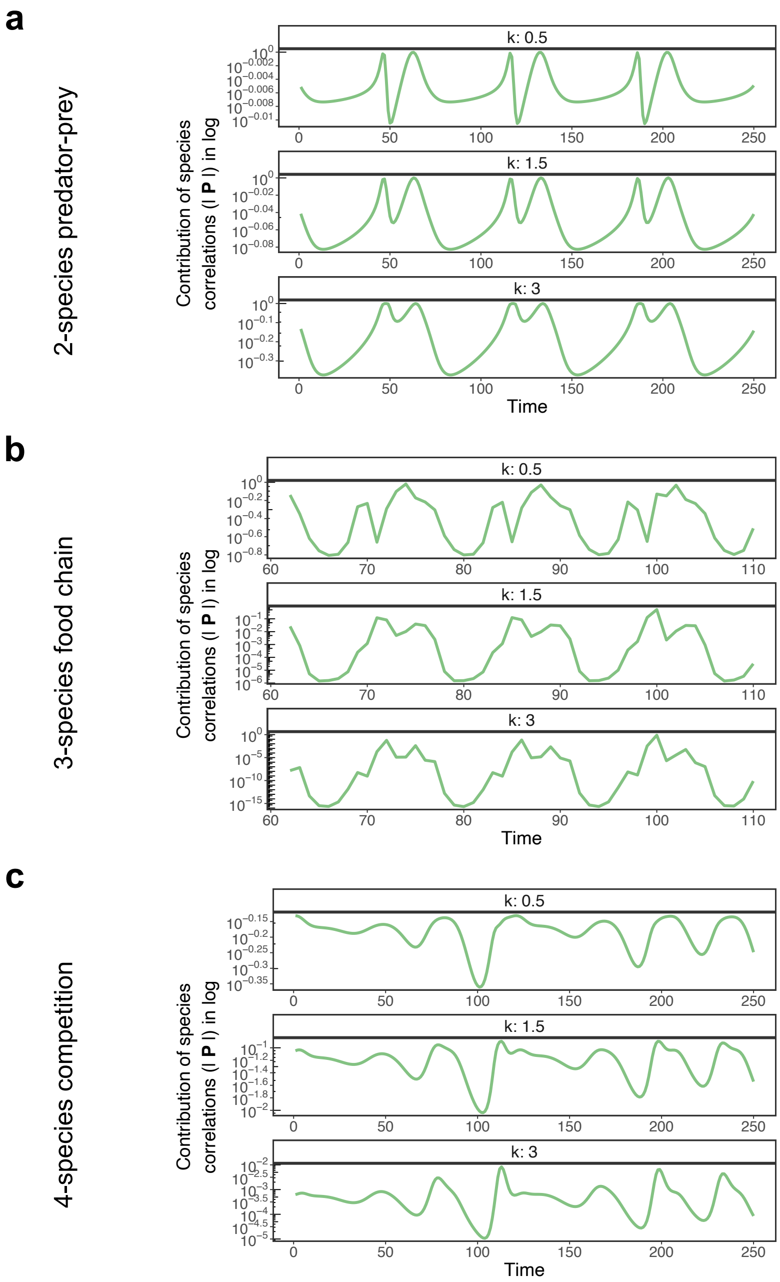


**Figure S5. Contribution of species correlations ($\log |\mathbf{P}|$) shows the same qualitative pattern over time for different values of $k$. a-c**, Each plot shows $\log |\mathbf{P}|$ over time computed using the analytical Jacobian matrix $\mathbf{J}$ and a given value of the time step $k$ (see *Materials and Methods* section in the main text). Note that the plots with $k = 3$ for the 2-species predator-prey (**a**) and the 4-species competition (**c**) models as well as the plot with $k = 0.5$ for the 3-species food chain model are identical to the center panels in Fig. 3 in the main text.

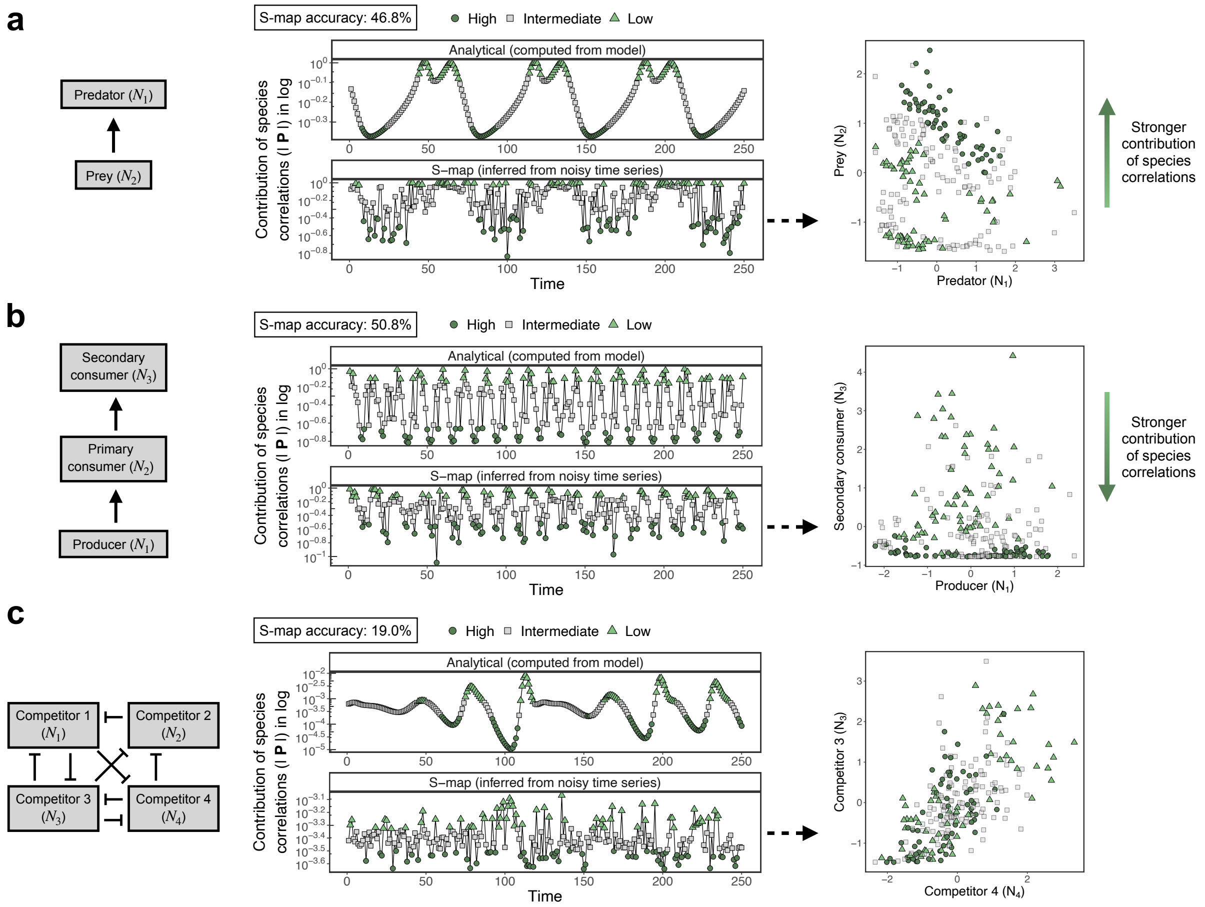


**Figure S6. State-dependent impact of species correlated responses can be inferred directly from time series with a high amount of noise in species abundances.** This figure is similar to Fig. 4 in the main text, but here we performed the S-map to infer the Jacobian matrix $\mathbf{J}$ using time series with 20% instead of 10% of observational noise (see *Materials and Methods* section in the main text). Note that the top center panels in **a**, **b**, and **c** (i.e., $\log|\mathbf{P}|$ computed analytically from model) are identical to the corresponding panels in Fig. 4.

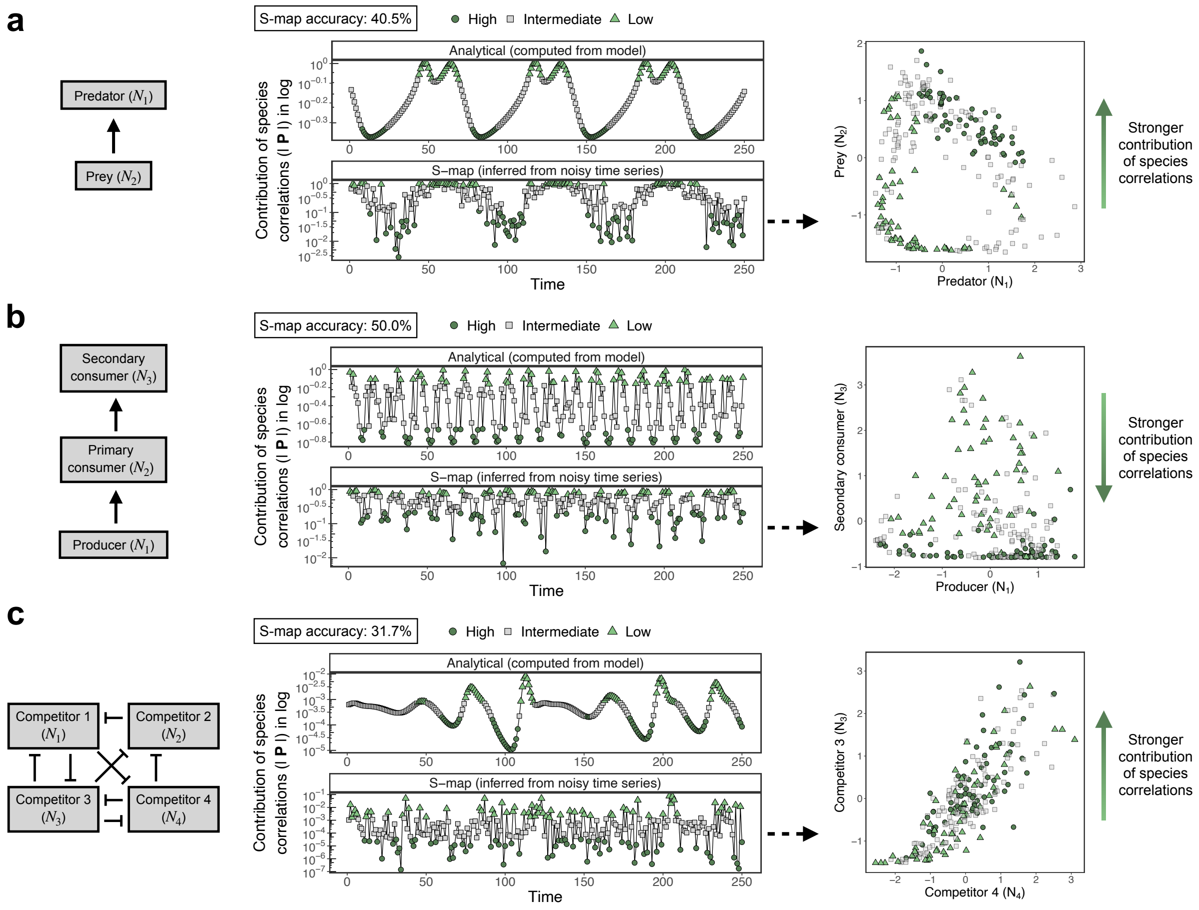


**Figure S7. State-dependent impact of species correlated responses can be inferred directly from time series with noise in $\mathbf{\Sigma}_t$ and $k$.** This figure is similar to Fig. 4 in the main text, but here we added noise to the covariance matrix of perturbations at time $t$ ($\mathbf{\Sigma}_t$) and to the time step $k$ before using $\mathbf{\Sigma}_t$ and $k$ to compute the covariance matrix of perturbations at time $t+k$ ($\mathbf{\Sigma}$, see *Materials and Methods* section in the main text) with the S-map. Specifically, for each time series point ($\mathbf{N}(t)$), we added 20% of Gaussian noise to the diagonal elements of $\mathbf{\Sigma}_t$ (i.e., perturbation variances) and to $k$ and then used the noisy $\mathbf{\Sigma}_t$ and $k$ to compute $\mathbf{\Sigma}$, which was in turn used to compute $\log|\mathbf{P}|$. Note that we only added noise when computing $\mathbf{\Sigma}$ with the S-map and the top center panels in **a**, **b**, and **c** (i.e., $\log|\mathbf{P}|$ computed analytically from model) are identical to the corresponding panels in Fig. 4. Also note that there is also 10% of observational noise in species abundances in addition to noise in $\mathbf{\Sigma}_t$ and $k$.

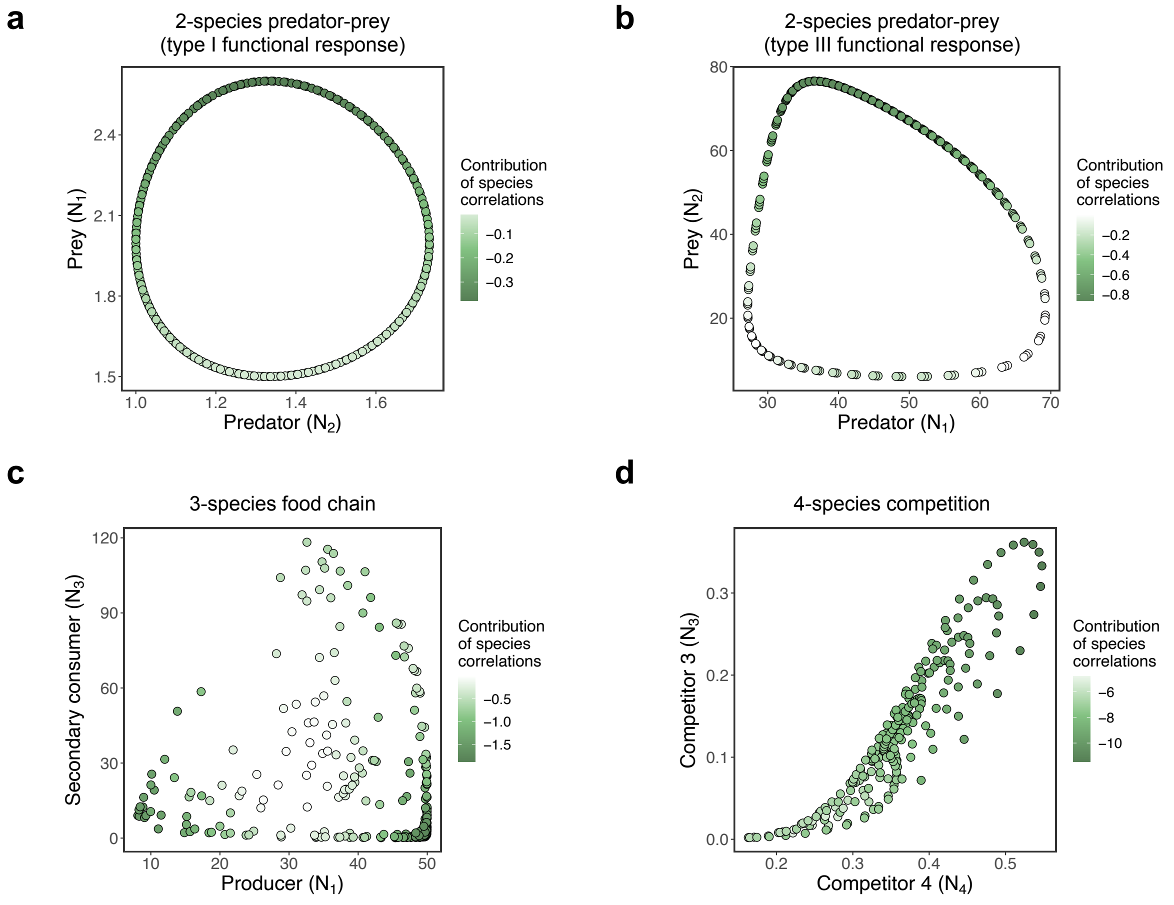


**Figure S8. Contribution of species correlations ($\log|\mathbf{P}|$) across state space under four different population dynamics models. a-d**, Each plot shows the attractor in state space corresponding to a given synthetic time series generated by the indicated population dynamics model. Each abundance state $\mathbf{N}(t)$ (i.e., each point) is colored according to $\log|\mathbf{P}|$ computed analytically from the model (see *Materials and Methods* in the main text). Note that panel **a** depicts a Lotka-Volterra predator-prey model (not shown in the main text) generated using the following parameters in equation [4] in the main text: $S = 2$, $r_1 = 0.2$, $r_2 = -0.2$, $a_{11} = 0$, $a_{12} = -0.15$, $a_{21} = 0.1$, and $a_{22} = 0$. Importantly, note that for both 2-species predator-prey models (**a** and **b**), $\log|\mathbf{P}|$ is higher when the prey abundance is higher.

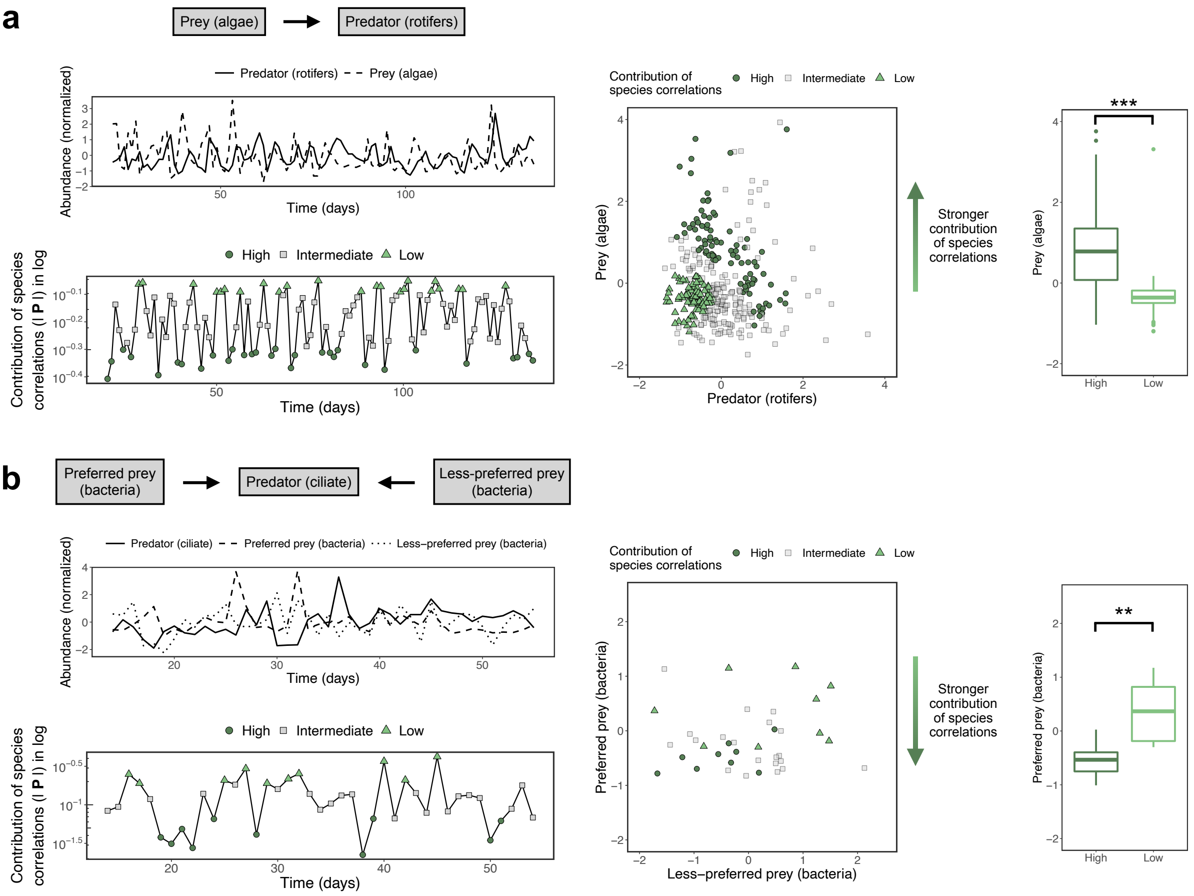


**Figure S9. Impact of species correlated responses depends on prey abundance for two experimental communities using time step $k = 3$ instead of $k = 1$.** This figure is similar to Fig. 5 in the main text, but here we use $k = 3$ instead of $k = 1$ to compute the covariance matrix $\mathbf{\Sigma}$, which is in turn used to compute the contribution of species correlations ($\log|\mathbf{P}|$). Note that results are qualitatively the same as in Fig. 5. For the 2-species community (**a**), prey abundance is higher when the contribution of species correlations is high than when it is low (two sample t-test: $t(130.52) = 9.91$, $p < 0.0001$). For the 3-species community, the abundance of the preferred prey is lower when the contribution of species correlations is high than when it is low (two sample t-test: $t(10.12) = -3.14$, $p = 0.01$).
